# A Precision Functional Atlas of Network Probabilities and Individual-Specific Network Topography

**DOI:** 10.1101/2022.01.12.475422

**Authors:** Robert J.M. Hermosillo, Lucille A. Moore, Eric Fezcko, Ally Dworetsky, Adam Pines, Gregory Conan, Michael A. Mooney, Anita Randolph, Babatunde Adeyemo, Eric Earl, Anders Perrone, Cristian Morales Carrasco, Johnny Uriarte-Lopez, Kathy Snider, Olivia Doyle, Michaela Cordova, Bonnie J. Nagel, Sarah W. Feldstein Ewing, Theodore Satterthwaite, Nico Dosenbach, Caterina Gratton, Steven Petersen, Óscar Miranda-Domínguez, Damien A. Fair

## Abstract

The brain is organized into a broad set of functional neural networks. These networks and their various characteristics have been described and scrutinized through *in vivo* resting state functional magnetic resonance imaging (rs-fMRI). While the basic properties of networks are generally similar between healthy individuals, there is vast variability in the precise topography across the population. These individual differences are often lost in population studies due to population averaging which assumes topographical uniformity. We leveraged precision brain mapping methods to establish a new open-source, method-flexible set of precision functional network atlases: the Masonic Institute for the Developing Brain (MIDB) Precision Brain Atlas. Using participants from the Adolescent Brain Cognitive Development (ABCD) study, single subject precision network maps were generated with two supervised network-matching procedures (template matching and non-negative matrix factorization), an overlapping template matching method for identifying integration zones, as well as an unsupervised community detection algorithm (Infomap). From these individualized maps we also generated probabilistic network maps and integration zones for two demographically-matched groups of n∼3000 each. We demonstrate high reproducibility between groups (Pearson’s r >0.999) and between methods (r=0.96), revealing both regions of high invariance and high variability. Compared to using parcellations based on groups averages, the MIDB Precision Brain Atlas allowed us to derive a set of brain regions that are largely invariant in network topography across populations, which provides more reproducible statistical maps of executive function in brain-wide associations. We also explore an example use case for probabilistic maps, highlighting their potential for use in targeted neuromodulation. The MIDB Precision Brain Atlas is expandable to alternative datasets and methods and is provided open-source with an online web interface to encourage the scientific community to experiment with probabilistic atlases and individual-specific topographies to more precisely relate network phenomenon to functional organization of the human brain.

## INTRODUCTION

Over the last decades, there have been several attempts to generate representations that delineate homogenous functional brain areas into parcellations or networks for use in non-invasive neuroimaging. These efforts have led to a series of structure and function-based parcellations that investigators use for various types of Brain-Wide Association Studies (BWAS). These regional parcellations of network descriptions are often based on group-averaged data (Fan et al., 2016; Glasser and Van Essen, 2011; Glasser et al., 2016; Gordon et al., 2014, 2016; Power et al., 2011; Yeo et al., 2011). However, there is significant inter-subject variability in topography on the macroscopic scale (Brodmann, 1909; Churchland and Sejnowski, 1988; Cui et al., 2020; von Economo and Koskinas, 1925; Glasser et al., 2016; Gordon et al., 2017a; Gratton et al., 2018a, 2020; Huth et al., 2016; Laumann et al., 2015; Rajkowska and Goldman-Rakic, 1995; Wang et al., 2015), which may limit power in BWAS (Feczko et al., 2021; Marek et al., 2020), or applicability of these parcellations to person-specific interventions (Cash et al., 2020, 2021a, 2021b; Fox et al., 2012a, 2012b).

Until recently, limited investigations have attempted to clearly describe individual variation of network-level topographical organization. While there is some degree of shared patterns of network organization among healthy populations, it is clear that large-scale brain networks show specific deviations from the group organization that are relatively stable (Gordon et al., 2017a; Laumann et al., 2015; Seitzman et al., 2019). Laumann and colleagues were the first to examine the personalized network structure using data from the MyConnectome Project (Laumann et al., 2015). Building on prior work using data-driven community detection to identify separable networks in the brain (Fair et al., 2009; Power et al., 2011), the authors precisely mapped these networks in an individual from whom they had collected more than 14 hours of resting state data (Laumann et al., 2015; Poldrack et al., 2015). These experiments revealed that while individuals have broadly similar networks to those identified in group averages, specific aspects of the topographical organization of these systems are highly unique.

Several studies have now shown that precisely mapping an individual’s brain may require upwards of 40-60 minutes of resting state data (Gordon et al., 2017a; Laumann et al., 2015). However, the collection of 40-60 minutes worth of data per participant is a burden to the participant and expensive for the investigative team and therefore creates limitations for widespread adoption. Extended collection of resting data creates additional obstacles for studies in childhood development and disease research, where a resting state session is typically limited to shorter durations. For example, the ABCD study was designed to collect resting state and task fMRI data in participants representative of the United States population at baseline years of 9-10 years old and biennially for 10 years (currently 11,987 participants enrolled). The goal of the ABCD Study is to determine biological and environmental factors that impact brain function by creating collaborative data resources that model the human brain development in childhood (Casey et al., 2018; Volkow et al., 2018). Though ABCD will provide an impressive resource for describing individual variation in network organization over time, “only” 20 minutes of resting state data is collected per participant, which may reduce the ability to maximize the precision of the individualized connectome across all participants. However, the shorter resting state data set is still valuable for precision mapping using new ‘supervised’ methods (Dworetsky et al., 2020; Gordon et al., 2017b) that create individual-specific networks that may only be marginally less precise. Furthermore, as task activity only adds a relatively small amount of variance to global resting-state brain organization (Gratton et al., 2018a), the additional task fMRI data (40 minutes) per participant can be used to generate individual-specific networks using similar amounts of data as prior reports (Cui et al., 2020; Gordon et al., 2017a; Laumann et al., 2015). In addition, the combination of a relatively large sample from ABCD (N>10,000), with relatively long BOLD data collected from each participant, provides the unique opportunity to provide individual network topographies and to produce a probabilistic atlas of functional networks.

Historically, probabilistic atlases in neuroimaging have been structural, not functional. For example, the standard Montreal Neurological Institute (MNI) (Evans et al., 1993), and other freely-available brain structural atlases based on several hundred individual MRI scans (Fonov et al., 2011; Mazziotta et al., 2001; Wang et al., 2013) are regularly used for image-based registration (Avants et al., 2009). These procedures often use probabilistic weights to attempt to correctly delineate anatomical structures, such as the whole cortex (Wang et al., 2013), amygdala (Tyszka et al., 2016), basal ganglia (Keuken and Forstmann, 2015), and brain stem nuclei (Pauli et al., 2018). Probabilistic volumes for subcortical structures are also often associated with these atlases, providing probabilistic-based ROIs. Just as these methods have vastly improved standard structural registration and segmentation, functional probabilistic maps may also be leveraged to create individual-specific functional mappings in group-level studies that would normally lack sufficient amounts of data for individual-specific mapping.

In addition, where resting state data is unavailable or researchers are unable to leverage community-detection techniques for precision brain mapping, these functional atlases may also provide unprecedented confidence in neuronavigation for targeted brain stimulation based on function and not simply anatomical landmarks.

Building on recent reports using probabilistic map approaches to resting state functional connectivity (rs-fcMRI) (Dworetsky et al., 2020), we implement various methods of network identification: Infomap (Gordon et al., 2017a; Laumann et al., 2015; Rosvall and Bergstrom, 2007, 2008), template matching (TM) (Dworetsky et al., 2020; Gordon et al., 2017b), non-negative matrix factorization (Cui et al., 2020; Li et al., 2017), and a novel overlapping network method to generate individual-specific network mappings, along with a population network probability atlases from resting-state fMRI data from the ABCD study. The highly reproducible probabilistic atlases enable derivation of ROI sets that reflect the variation of brain topography of individuals.

In the current report, we release the MIDB Precision Brain Atlas that includes: individual-specific networks, population probabilistic maps, individual integrative zones, and population integrative zones. We also describe this comprehensive MIDB Precision Brain Atlas, and we encourage others to contribute their own individualized networks and probabilistic maps of functional neural networks to the MIDB Precision Brain Atlas. In addition to the ABCD probabilistic maps provided upon the initial MIDB Precision Brain Atlas release, we are sharing additional probabilistic maps generated by Dworestky and colleagues (Dworetsky et al., 2020) from a Washington University dataset (Power et al., 2012), a Dartmouth dataset (Gordon et al., 2016), the Midnight Scan Club (MSC) dataset (Gordon et al., 2017a), and the Human Connectome Project (HCP) dataset (Van Essen et al., 2012). Furthermore, the resource also includes a user-friendly downloader tool with an adjustable thresholding of network assignment and functional integration zones. As a resource available to the scientific community, the MIDB Precision Brain Atlas will enable systematic investigation of the contributions of network topography and network-network interaction to human cognition and behavior.

## RESULTS

### ABCD Cohort Demographics

The ABCD data were divided into 2 large cohorts (discovery Cohort ABCD-1 n=5786, replication Cohort ABCD-2 n=5786), and 1 smaller test cohort (Cohort ABCD-3 n=300), matched on multiple demographics (see Supplementary Table 1) (i.e. the ABCD reproducible matched samples [ARMS] from the ABCD BIDS Community Collection [ABCC](Feczko et al., 2021)). From these initial groups, participants with at least 10 minutes of low motion data were chosen to test replication (Group 1 n=2995, Group 2 n=3111; see Supplementary material for analysis of required minutes) based on a framewise displacement (FD)<0.2, which retained similar proportional demographics to that of the full cohort (Table 1; see and supplementary Figure S1). Group 3 (n=164 with available processed MRI data) was used to build the network templates for the template matching (TM) procedure described below. Groups 1 and 2 were test groups used to validate the community detection methods.

**Table 1:** Demographics table – Subjects with at least 10 minutes of resting state data. ( from resource paper)

|  |  |  |  |
| --- | --- | --- | --- |
| Variable | Group1 (N=2995) | Group2 (N=3111) | Group3 (N=161) |
|  | mean (sd) | mean (sd) | mean (sd) |
| Age (in months) | 119.64 (7.48) | 119.75 (7.47) | 118.37 (7.73) |
| Grade level | 4.27 (0.78) | 4.27 (0.78) | 4.20 (0.76) |
| Highest parent education | 17.38 (2.85) | 17.34 (2.46) | 16.83 (2.90) |
| Combined income (in thousands). | 7.51 (2.24) | 7.46 (2.24) | 7.08 (2.35) |
| categorical | Group1 (N=2995) | Group2 (N=3111) | Group3 (N=161) |
|  | count (%) | count (%) | count (%) |
| # Female* | 1411 (47.10) | 1544 (49.66) | 78 (48.45) |
| Anesthesia exposure | 966 (32.2) | 1005 (32.3) | 42 (26.1) |
| Right handed | 2401 (80.2) | 2525 (81.2) | 136 (84.5) |
| Race/Eth |  |  |  |
| White | 2399 (80.10) | 2460 (79.07) | 106 (65.84) |
| Black | 539 (18.00) | 556 (17.87) | 30 (18.63) |
| AlAK** | 94 (3.14) | 99 (3.18) | 4 (2.48) |
| NHPI | 19 (0.63) | 18 (0.58) | 0(0) |
| Asian | 62 (2.07) | 62 (1.99) | 6 (0.373) |
| Other | 143 (4.77) | 166 (5.34) | 21 (13.04) |
| Unknown | 23 (0.77) | 34 (1.09) | 4 (2.48) |
| Latinx | 544 (18.16) | 564 (18.13) | 28 (17.39) |
| site | Group1 (N=2995) | Group2 (N=3111) | Group3 (N=161) |
|  | count (%) | count (%) | count (%) |
| 1 | 41 (1.37) | 49 (1.58) | 4 (2.48) |
| 2 | 212 (7.08) | 205 (6.59) | 9 (5.59) |
| 3 | 199 (6.64) | 213 (6.85) | 5 (3.11) |
| 4 | 205 (6.84) | 182 (5.85) | 9 (5.59) |
| 5 | 103 (3.44) | 112 (3.60) | 12 (7.45) |
| 6 | 170 (5.68) | 170 (5.46) | 22 (13.66) |
| 7 | 91 (3.04) | 84 (2.70) | 4 (2.48) |
| 8 | 62 (2.07) | 88 (2.83) | 3 (1.86) |
| 9 | 122 (4.07) | 112 (3.60) | 6 (3.73) |
| 10 | 128 (4.27) | 146 (4.69) | 5 (3.11) |
| 11 | 133 (4.44) | 122 (3.92) | 4 (2.48) |
| 12 | 42 (1.40) | 57 (1.83) | 3 (1.86) |
| 13 | 157 (5.24) | 145 (4.66) | 5 (3.11) |
| 14 | 189 (6.31) | 194 (6.24) | 11 (6.83) |
| 15 | 89 (2.97) | 95 (3.05) | 8 (4.97) |
| 16 | 380 (12.69) | 404 (12.99) | 10 (6.21) |
| 17 | 97 (3.24) | 121 (3.89) | 8 (4.97) |
| 18 | 88 (2.94) | 86 (2.76) | 2 (1.24) |
| 19 | 124 (4.14) | 115 (3.70) | 12 (7.45) |
| 20 | 183 (6.11) | 212 (6.81) | 10 (6.21) |
| 21 | 180 (6.01) | 199 (6.40) | 9 (5.59) |

### Individual-specific network mapping through multiple community detection techniques

We first sought to establish that each method produces consistent within-subject networks by demonstrating that a given subject is distinguishable from the group. Individual-specific networks were successfully mapped using the following methods: Infomap (IM) (Gordon et al., 2017a; Laumann et al., 2015; Rosvall and Bergstrom, 2007, 2008) and TM (Dworetsky et al., 2020; Gordon et al., 2017b). For all participants in Groups 1 and 2, we created individual-specific network maps by generating dense connectivity matrices (91282 x 91282 grayordinates) from exactly 10 minutes of resting state data randomly sampled from the full length of data below a framewise displacement (FD) threshold of 0.2 mm (Supplementary Figure S3A, see Methods). This allowed for direct participant-to-participant comparison despite differences in movement characteristics within the MRI scanner between participants. Identical matrices were used for each individual as inputs for both the Infomap and TM procedures (see below).

The Infomap algorithm models flow between nodes of a graph by using information theory to map networks or modules. Specifically, the algorithm implements a random walk strategy using the connection weights, in an attempt to minimize the number of bits (using Huffman coding) necessary to identify the module (i.e. network) to which each node (i.e grayordinate) belongs. Each connectivity matrix was thresholded to discrete percentiles of connections (or edges), then infomap was used to identify community structure (see Methods) at each of these thresholds. Lastly, we generated a consensus across the edge densities to: 1) ensure that similar communities are identified among the groups, 2) ensure that distinct communities at smaller percentiles, are accurately assigned to larger networks, and 3) provide brain coverage, as in previous work (Gordon et al., 2017a).

The TM algorithm, assigns each grayordinate to a network by comparing the whole-brain connectivity of the grayordinate to a series of network templates observed in the group (Dworetsky et al., 2020; Gordon et al., 2016), a method developed by Gordon and colleagues (Gordon et al., 2017b). Supplementary Figure S3 shows the technique used to establish individual-specific networks using TM. Using Group 3 participants, templates were generated for each network. This was done by using network templates previously identified with Infomap with an average dense connectivity matrix from 120 adult participants (Gordon et al., 2017a; Laumann et al., 2015; Rosvall and Bergstrom, 2007, 2008). We then conducted a seed-based correlation whereby the motion-censored (see below) resting state data for each grayordinate is correlated with the average resting state signal for the respective network. To perform TM, we used the network templates generated with Group 3 to measure the extent to which the whole-brain connectivity of each grayordinate resembles the connectivity pattern of the template network for each subject in Groups 1 and 2 (Supplementary Figure S3A) (using eta^2^). Supplementary Figure S4 shows the network templates that were used, which correspond closely with networks that have been previously identified within the literature (Gordon et al., 2017b; Gratton et al., 2018a; Harrison et al., 2015; Li et al., 2017; Marek et al., 2018, 2019). All of these individual-specific maps are available on the NDA via the ABCC (Feczko et al., 2020, 2021).

### Network mapping methods demonstrate high intra-subject reliability

In order to establish that these methods can reliably generate individual-specific network maps using limited amounts of data (i.e. 10 minutes of motion-free data), we used split-half reliability analysis to demonstrate that similar network maps are generated when using the first vs. second half of a subject’s timeseries (Figure 1). We conducted split-half reliability analyses for each method in 10 participants from Group 1 who had the longest duration of low-motion quality data, exceeding 20 minutes. To assess network reliability, we used normalized mutual information (NMI). We compared the topographic similarity of network maps generated within-subject (intra-subject; first half of data vs. second half of data) to network maps generated between different subjects (inter-subject). The distribution of the NMI between intra-subject halves was compared against a null distribution of the NMI between inter-subject halves (Figure 1B). For both TM and Infomap, intra-subject NMI was significantly higher than inter-subject NMI (TM: t(9.31)=11.87, p=3.1079 x 10^-7^; IM: t(9.607)=9.049, p=2.6109 x 10^-6^; unequal variance assumed, one tailed). Comparing methods, TM displayed significantly higher similarity both between halves of data from the same subject (mean NMI for TM=0.421; IM=0.370, t(18)=2.951, p=0.009) and different subjects (t(358)=16.0315, p=5.60 x 10^-44^, equal variance assumed, two-tailed, Figure 1B vs C, gold bars). TM had a higher between-group similarity compared to infomap when we used group-averaged connectivity matrices (Supplementary Figure S6). Overall, despite intrasubject variability, these data highlight that networks generated by both methods are highly specific to each individual.

**Figure 1.**
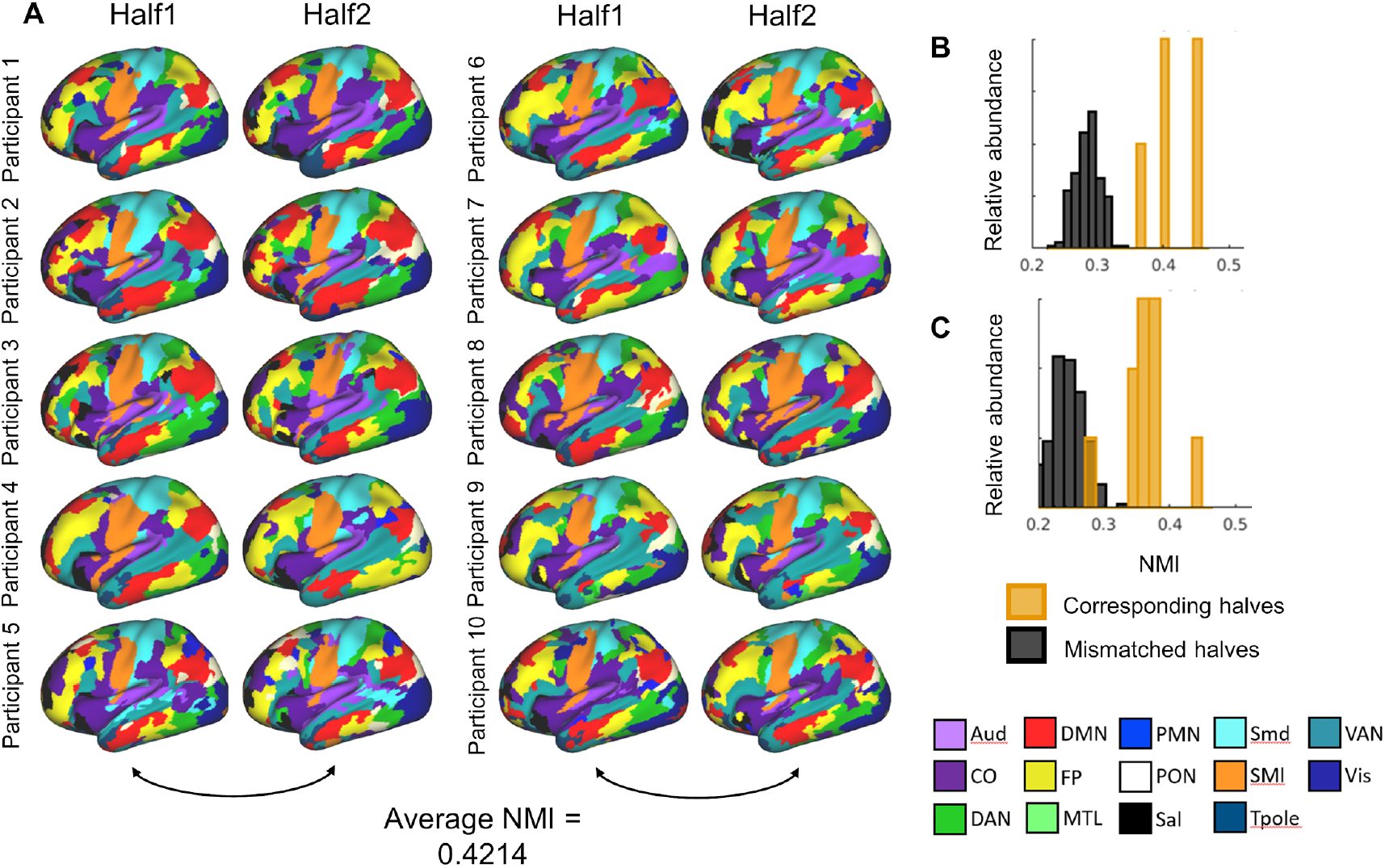
Example of precision maps of ABCD participants using template matching. **A)** Example of networks determined by the template matching procedure for subjects with at least 20 minutes of low motion resting state data. Resting state time series were split in half and networks were obtained for each half (n=10). Only the left hemisphere is shown for visualization purposes, but networks were also identified in the right hemisphere, subcortex, and the cerebellum. **B)** The normalized mutual information (NMI) was calculated between participants’ own halves (gold bars) and others in the split half group (grey bars) using template matching. **C)** We also generated network maps using the Infomap procedure and performed an identical NMI comparison (maps not shown).

### Minutes of resting state data required to produce reasonable specificity for TM

Next, we tested the minimum necessary resting state data (in minutes of low-motion data) required to produce individual-specific network maps. Using publicly available resting state data from the Midnight Scan Club (MSC) (Gordon et al., 2017a), we conducted an analysis similar to the split halves reliability analysis conducted with ABCD. However, to perform the split halves reliability analysis for networks generated with the TM procedure, split halves were made using interleaved concatenated sessions (see supplement for further details). Individual-specific networks were generated from 1, 2, 3, 4, 5, 10, 15, and 20 randomly sampled non-contiguous minutes of low-motion data (10 times each) from one half of each subject‘s data and compared to networks generated from the second half of data using NMI. We demonstrate that individual-specific maps produced by TM, even with relatively few minutes of low-motion data, reliably resemble the individual-specific network maps observed with 10-minutes of low-motion data (See Supplementary Figure S5). It should be noted that randomly-sampled data from longer duration acquisitions likely improves reliability (increase in correlation up to 0.04) due to the diminished influence of autocorrelation in the time series data (Laumann et al., 2015).

### Calculating the probability of observing a given network at each grayordinate

Next, we illustrated the extent to which each grayoridinate participates in each network across both ABCD Groups 1 and 2. Using the individual-specific mapping methods described above, we generated probabilistic maps in both ABCD Groups 1 and 2 to highlight replicable network probabilities between the groups. Individual-specific maps were generated for each participant within Groups ABCD-1 and 2, then the probability of network observation was calculated for each grayordinate (Figures 2A and 2D). To test replication between groups and methods, we correlated non-zero values of probabilistic maps. For example, the frontoparietal network (Figures 2A and 2E) show remarkable replicability (r=0.9996, Figure 2I) between groups, even with respect to functional asymmetries. Note how the frontoparietal representation in the dorsolateral prefrontal cortex (DLPFC) in the right hemisphere compared to the left is clearly present in both groups. Additionally, these probabilistic maps highlight the discrete nuclei of the cerebellum that communicate heavily with the frontoparietal network (Figure 2K), and the location of these nuclei highly spatially correspond with frontoparietal clusters that have been previously observed in the cerebellum (Marek et al., 2018).

**Figure 2.**
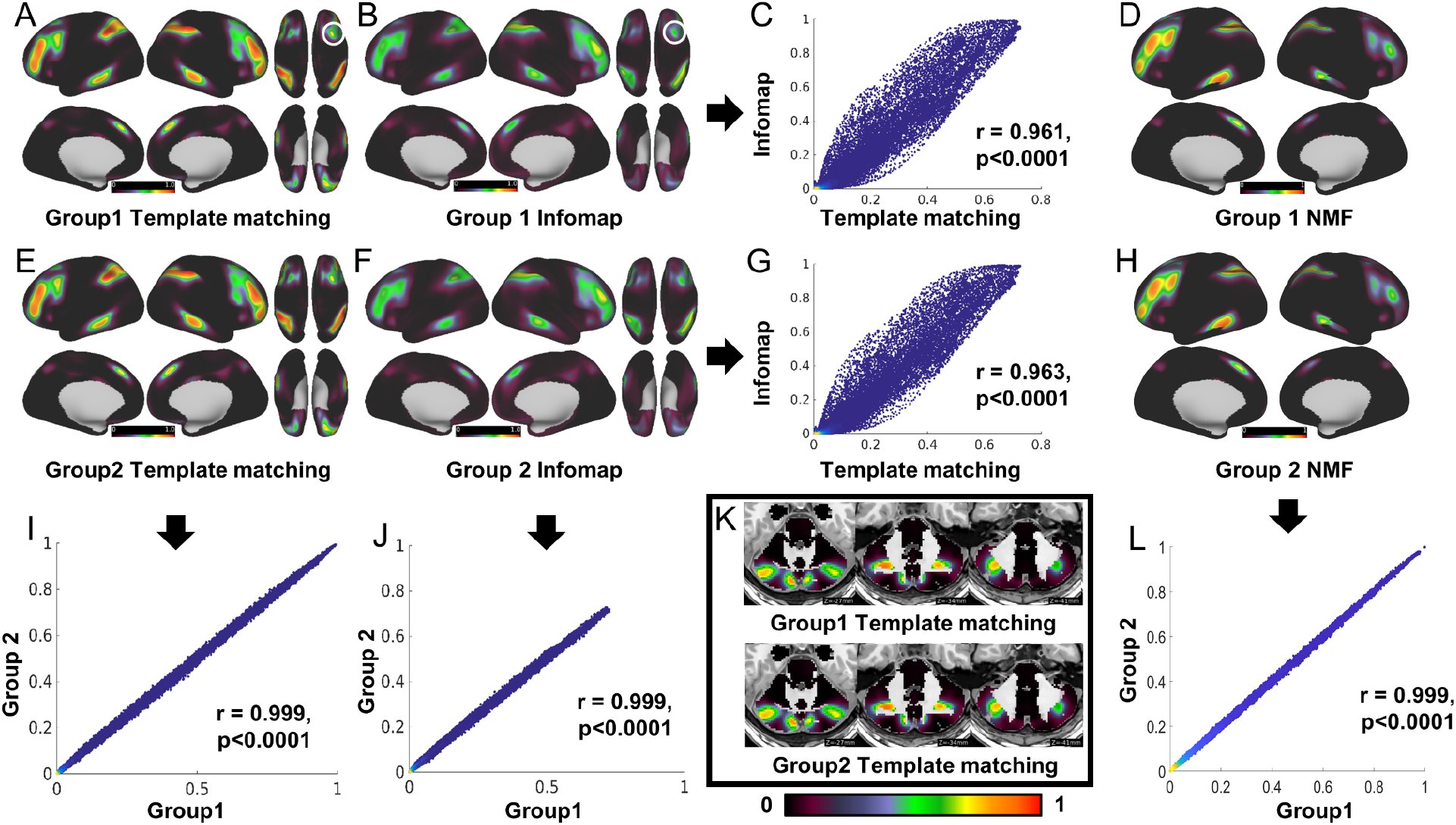
Example of Network probability. An example of Network probability for the frontoparietal network using template matching (**A and E**), infomap (**B and F**) (surface only), and NMF (**D and H**) (surface only) procedures with single network assignment. **I**, **J**, and **L** show the between-group correlation for template matching and Infomap respectively. **C** and **G** show the correlation between methods for Groups 1 and 2 respectively. For additional probability maps, see Supplementary Figure S1. **K)** Network probabilistic map for the frontoparietal network within the cerebellum (Infomap not shown). White circles in **A** and **B** highlight similar probabilistic functional asymmetries in the supplementary motor area (SMA) across methods. Each circle in **C**, **G**, **I**, **J**, and **L** represents 1 grayordinate. The scale is identical (0-1 Pearson’s r) to that shown in the surface maps.

To confirm that the probabilistic network representations observed in Figure 2A and 2E are not simply the product of the TM method, we used the same data to generate probabilistic maps using a robust community detection method for large-scale neuroimaging data, Infomap (Figures 2B and 2F) (Gordon et al., 2017a; Laumann et al., 2015; Rosvall and Bergstrom, 2007, 2008), and NMF (Figures 2D and 2H) (Cui et al., 2020; Lee and Seung, 1999). We compared methods by correlating the probabilistic maps between Infomap and TM for ABCD-group1 (Figure 2C) and ABCD-group2 (Figure 2G) respectively. Cross-method correlation analysis between NMF and other methods was not performed due to the differing number of unique networks.

Nevertheless, NMF probabilistic maps demonstrate very high correlation between groups (Figure 2L, r=0.9996, p <0.0001). Probabilistic network topography remains highly conserved across methods (albeit overall probability is slightly reduced in Infomap), suggesting that the supervised method produces nearly identical networks to an unsupervised approach (frontoparietal network: non-zero correlation: Group1: TM to IM: r(91282)=0.951, p<0.0001; Group2: TM to IM: r(91282)=0.954, p<0.0001), even retaining the aforementioned asymmetries. Supplementary Table 2 provides the correlation between methods for the remaining networks (median between-method correlation : ABCD-Group 1=0.937; ABCD-Group 2=0.936). Visual comparisons for all networks are provided (Supplementary Figures S8). In addition, we generated probabilistic maps for each network, using 10 minutes of low-motion data from the cerebral cortex only (i.e excluding subcortical nuclei and cerebellum) (Supplementary Figure S9).

To assess the impact of including task data to generate probabilistic maps, the same between-method comparison was performed using concatenated rest and task data instead of rest alone (Supplementary Figure 10). Including task data provided an additional 40 minutes of data per subject.

We observed strong replication between groups (median between-method correlation: ABCD-Group 1=0.900; ABCD-Group 2=0.900), but crucially, the probability maps are nearly identical to networks generated from resting state data alone, despite differing amounts of data used to generate the maps (See Supplemental text). Figure S10 shows the probability map for the network using TM with single network assignment, similar to what was shown in Figure S9 except with task data included (see Methods). This suggests that the contribution of task-induced, activation-related, hemodynamic responses do not appreciably affect global network organization, which others have previously proposed (Buckner et al., 2009; Gratton et al., 2018a).

### Probabilistic–based ROIs improve reliability in brain-wide association studies (BWAS)

Recent evidence suggests that connectivity-based BWAS show limited predictive power when using whole-brain associations (Marek et al., 2020), therefore we wanted to test if omitting network topographies that are highly variable would improve group reliability when we only used commonly-observed network locations. Using the resting-state probabilistic maps, we generated a set of network labels to examine connectivity among brain regions that are highly homogenous across participants (Supplementary Figure S3D). Figure 3 shows the regional network composition and connectivity matrix across both ABCD groups at an 80% threshold (i.e. a consensus network map for which at least 80% of the subjects were assigned to a respective network). Using these regions of high consensus (Figure 3A), we produced a parcellated connectivity matrix for each participant. The strength of the within- and between-network connectivity for each cohort was calculated using the MIDB Probabilistic parcellation (Figure 3C and D) vs the Gordon parcellation (Figure 3 3E and F), one of the most widely used parcellation schemas which are based on a population average (Gordon et al., 2016; Schaefer et al., 2018). We observed a significant correlation between the average functional connectivity for each group (Figures 3C-F: Pearson’s r, upper triangle: TM: r(3.16 x 10^3^)=0.998, p=0); Gordon parcellation: r(6.17 x 10^4^)=0.996, p=0) ). Compared to Gordon parcellation based on group averages, the MIDB Probabilistic parcellation set provides increased within-network connectivity strength between the two group matrices (Figure 3G) (average within network connectivity: Group 1: Gordon: 0.3421±0.1467 TM: 0.5208±0.149, t(24)=-3.0801, p=0.0026; Group 2: Gordon: 0.3421±0.1467 TM: 0.5189±0.149, t(24)=-3.0402, p=0.0028). This increase in connectivity strength is likely due to only including regions with consistent network assignment across the population, and therefore have inhomogeneous connectivity.

**Figure 3.**
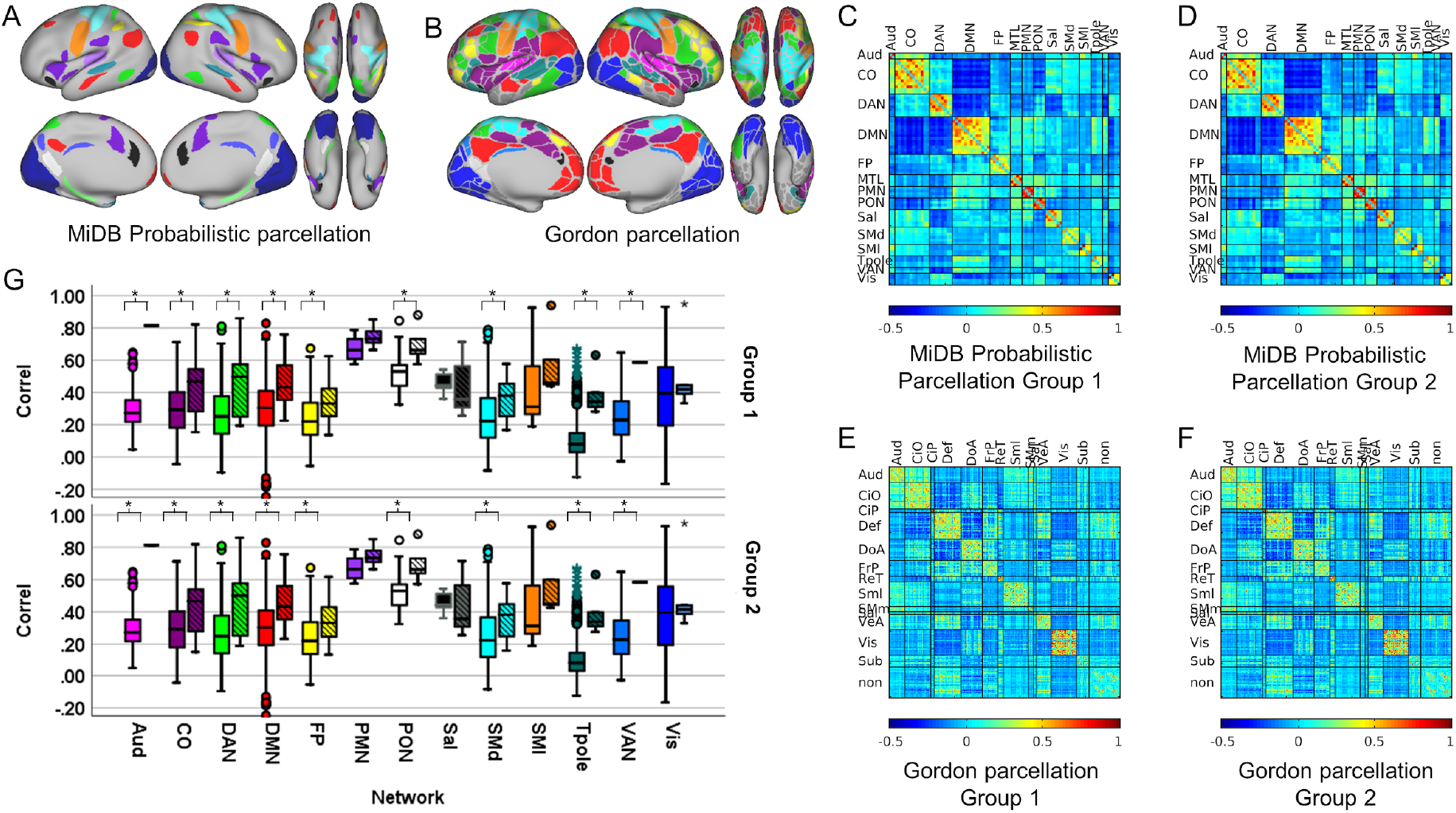
Comparing connectivity using probabilistic ROIs and Gordon ROIs. **A)** Probabilistic parcellation (80% probability of network consensus using the template matching). **B)** Gordon parcellation. Parcels are colored according to network assignment. Similar colors were used between parcellations where possible. **C-F)** Connectivity matrices were generated using the MiDB Probabilistic parcellations and the Gordon parcellation for groups 1 and 2. **G)** 9 of 13 shared networks, showed significantly higher within-network connectivity in the MiDB probabilistic parcellation compared to the Gordon parcellation. Open boxes=Gordon Parcellation; striped boxes=MiDB Probabilistic parcellation. * indicates alpha < 0.05.

In addition, we tested whether the probabilistic ROI sets provide additional reliability when performing brain-behavior correlations. Conventional ROIs sets that apply the same network assignment to the parcellation schema to all individuals have the potential to dilute the effects that specific brain regions have on behavior. For a given region of interest, several networks may include a given region, and furthermore, the same location may belong to different networks in any given individual (Figure 4A-B). A Bayesian probabilistic principal components analysis (BPPCA) was used to extract 3 cognitive traits from ARMS-1 and ARMS-2 separately reflecting general cognitive ability, executive function and working memory using previously published procedures (Feczko et al., 2021; Luciana et al., 2018; Thompson et al., 2019). We performed a subset reliability analysis using either the Gordon or MIDB probabilistic parcellation, sampling a random subset of Group 1 participants. We did this by correlating each element of the connectivity matrix from each subsample with each behavioral factor. We then correlated the brain-behavior correlations from each subsample in Group1 with the brain-behavior correlation using all Group 2 participants to serve as the “ground truth” (see Subset Reliability in Methods). When examining general cognitive ability, we found that using the MIDB probabilistic parcellation provided only a modest increase in reliability compared to using the Gordon ROIs at all sample sizes (Figure 4C), however, for the components of learning/memory and executive function, we observed an increase in reliability (Figures 4D-,F Cohen’s d with 1250 subjects: PC1=0.909, PC2=1.605,PC3=1.865). The MIDB probabilistic parcellation provided a substantial increase in reliability for associations with PC 1, 2,and 3. The maximum inter-group brain-behavior correlation that we observed with a subsample of 1250 subjects using the Gordon parcellation, could be observed with only 873 (PC1),702 subjects(PC2),and 675 subjects (PC3) Figure 4D-F.

**Figure 4.**
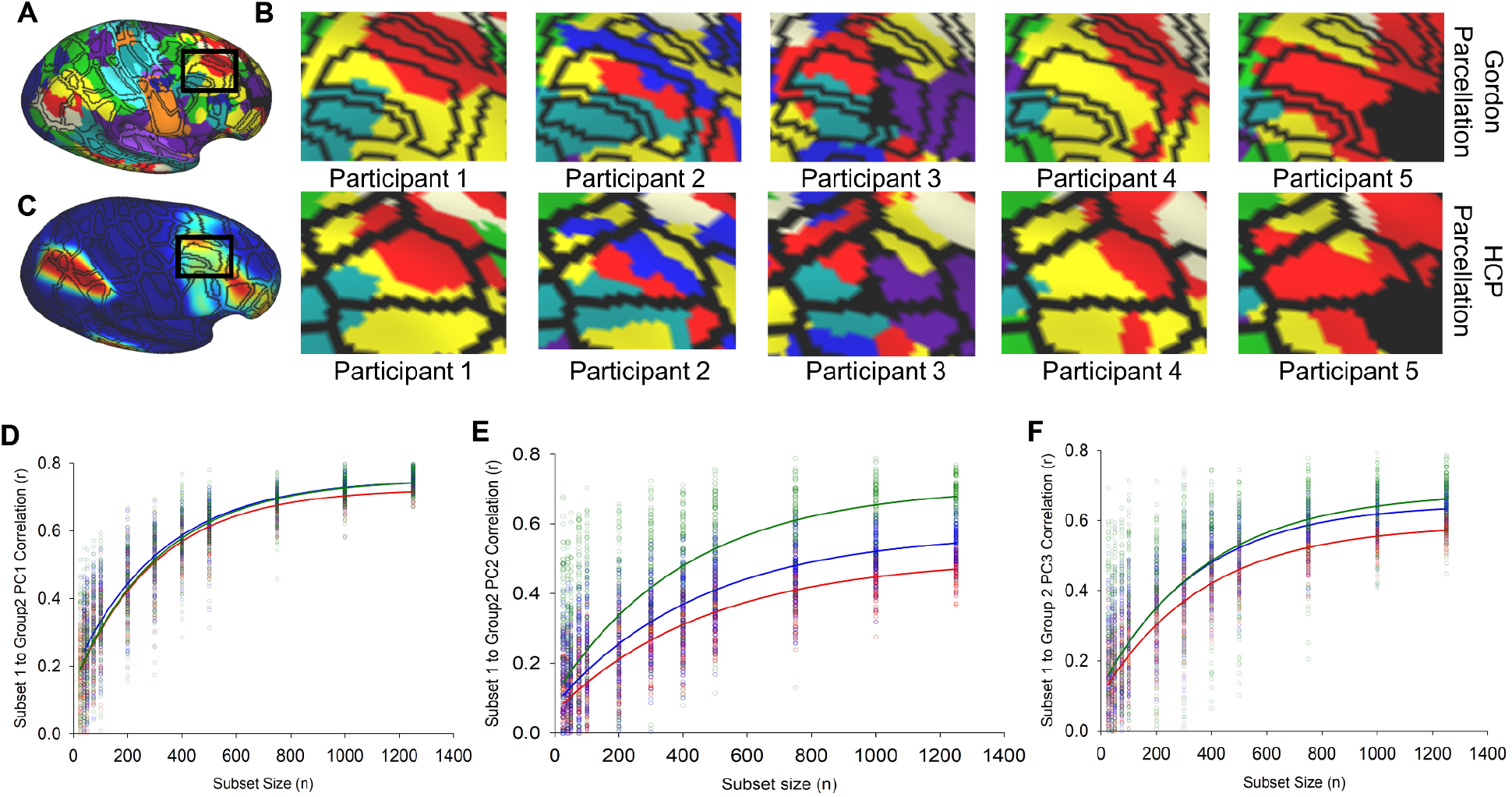
Neural networks have unique topographies that confound conventional ROI sets. **A)** The dorsolateral aspect of the frontal lobe demonstrates may belong to one of several potential networks. **B)** An example of 10 individuals’ neural networks with the Gordon Parcellation overlaid. Frontoparietal is shown as yellow. **C)** The Frontoparietal probabilistic map indicates inhomogeneity in network topography among the population. **D-F)** Subset reliability analysis showing that using the MiDB probabilistic parcellation improves signal-to-noise in group-level predictions relative to the Gordon Parcellation. Blue circles/lines indicate inter-group correlation for each random subset using the MIDB probabilistic parcellation. Red circles/lines indicate inter-group correlation for each random subset using the Gordon parcellation. Green circles/lines indicate inter-group correlation for each random subset using the Integration zone parcellation. Orange circles/lines indicate inter-group correlation for each random subset using 30 randomly the parcels from Gordon parcellation. Data were fitted with an exponential rise-to-maximum equation. Note: red and orange fitted curves are nearly identical which obscures visual discernment.

### Calculating network similarity at each grayordinate reveals an overlapping network structure

Most network connectivity studies to date assume that a given grayordinate (or voxel) participates in a single network. However, it has been suggested, and is likely, that some brain regions participate in multiple networks (Braga and Buckner, 2017), or demonstrate *nested*, or *hierarchical* structure that can be better described when allowing communities to overlap (Yang and Leskovec, 2013). For example, neurons that respond to multimodal stimuli likely participate in multiple networks (Andersen, 1997; Driver and Noesselt, 2008; Stein and Stanford, 2008). Therefore, we aimed to identify regions that belong to multiple communities by extending our TM procedure to allow networks to overlap (Gratton et al., 2018b; Greene et al., 2020; van den Heuvel and Sporns, 2013, 2019; Power et al., 2013a). We quantified the similarity of each grayordinate’s BOLD signal to observed networks by setting a data-driven threshold based on the observed local minima in the distribution of eta^2^ values calculated for each network used in TM (Figure 5). This technique allows us to detect secondary and tertiary (and so forth) networks that communicate with a particular grayordinate that would otherwise be missed by only identifying the primary network (see supplementary methods). For each network, we observed similar regions of high probabilistic similarity between ABCD groups1 and 2 for each of the networks measured (See Figure 6, Supplementary Figure S11) using the overlapping TM method.

**Figure 5.**
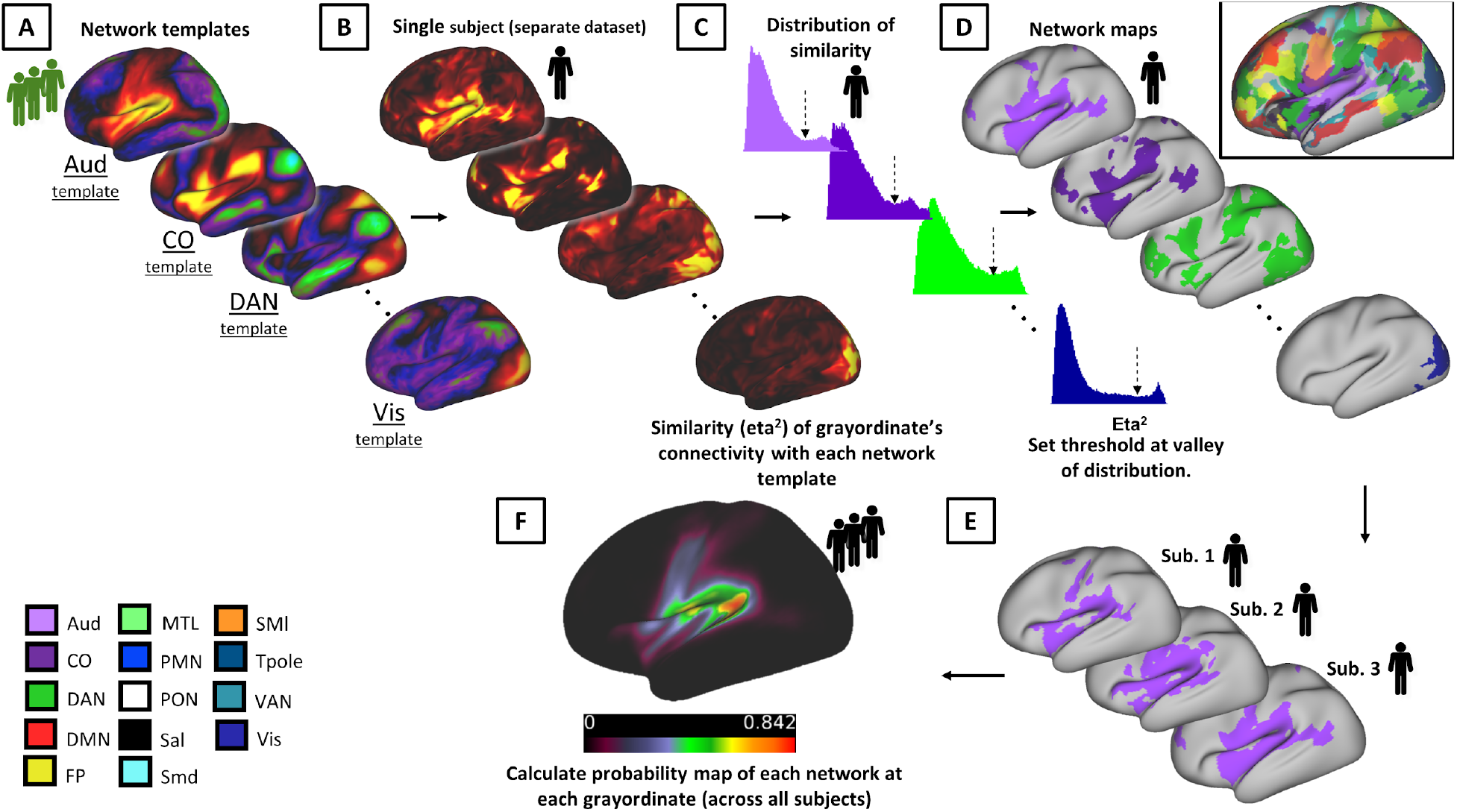
Method for detecting overlapping networks using template matching. **A)** A series of network templates were generated using an independent group of participants (ABCD-group 3). **B)** For each subject, the similarity at each grayordinate (using eta^2^) was calculated to each of the network templates shown in A. **C)** We set a threshold (dashed arrow) for each network, based on the observed local minimum between peaks of bimodal distribution of eta^2^. **D)** Grayordinates that had eta^2^ values that were above the threshold were then assigned that network label. All overlapping networks for an example subject are shown in the inset. **E,F)** After this procedure is performed for all subjects, we calculate a probabilistic map for each network (only the auditory network is shown).

**Figure 6.**
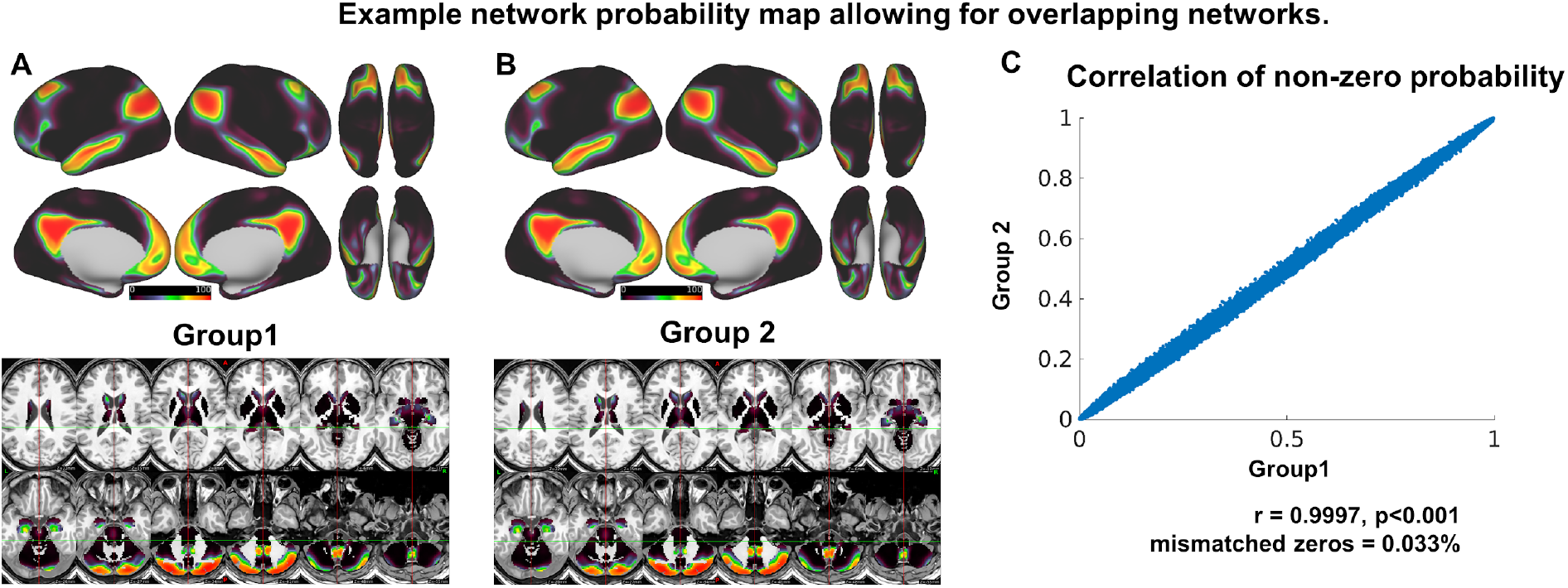
Network probability maps using overlapping networks. A, B) At each grayordinate, the probability of observing each network was calculated for ABCD group 1 and 2. Here the default mode network is shown as an example. **C)** The correlation of the probability maps depicted in A and B (excluding zeros). See Supplementary Figure S9 for additional networks.

### Integration Zones can be revealed by examining the number of overlapping networks

After generating the overlapping networks for an individual, we averaged the number of networks observed at each grayordinate across the group to examine the extent to which networks overlap in the population. Regions that demonstrate a high degree of overlap are thought to facilitate communication between networks (Gordon et al., 2018; Gratton et al., 2018b; van den Heuvel and Sporns, 2013, 2019; Power et al., 2013a).

Split-half reliability was calculated in the same manner with the 10 ABCD participants mentioned previously. Overlapping regions showed reliability within individuals (avg real NMI= 0.4847± 0.411 s.d.; null NMI=0.3287 ± 0.327 s.d., t(9.644) =11.783, uneq. var., p=4.84 x10^-7^) which was overall greater than using a single network assignment. In addition, we quantified the number of networks detected at each grayordinate. Figure 7A demonstrates that, within a given subject, some integrative zones can even show 8-10 networks converge in regions such as the posterior parietal cortex, precuneus, and posterior cerebellum, revealing a complex structure of internetwork communication.

**Figure 7.**
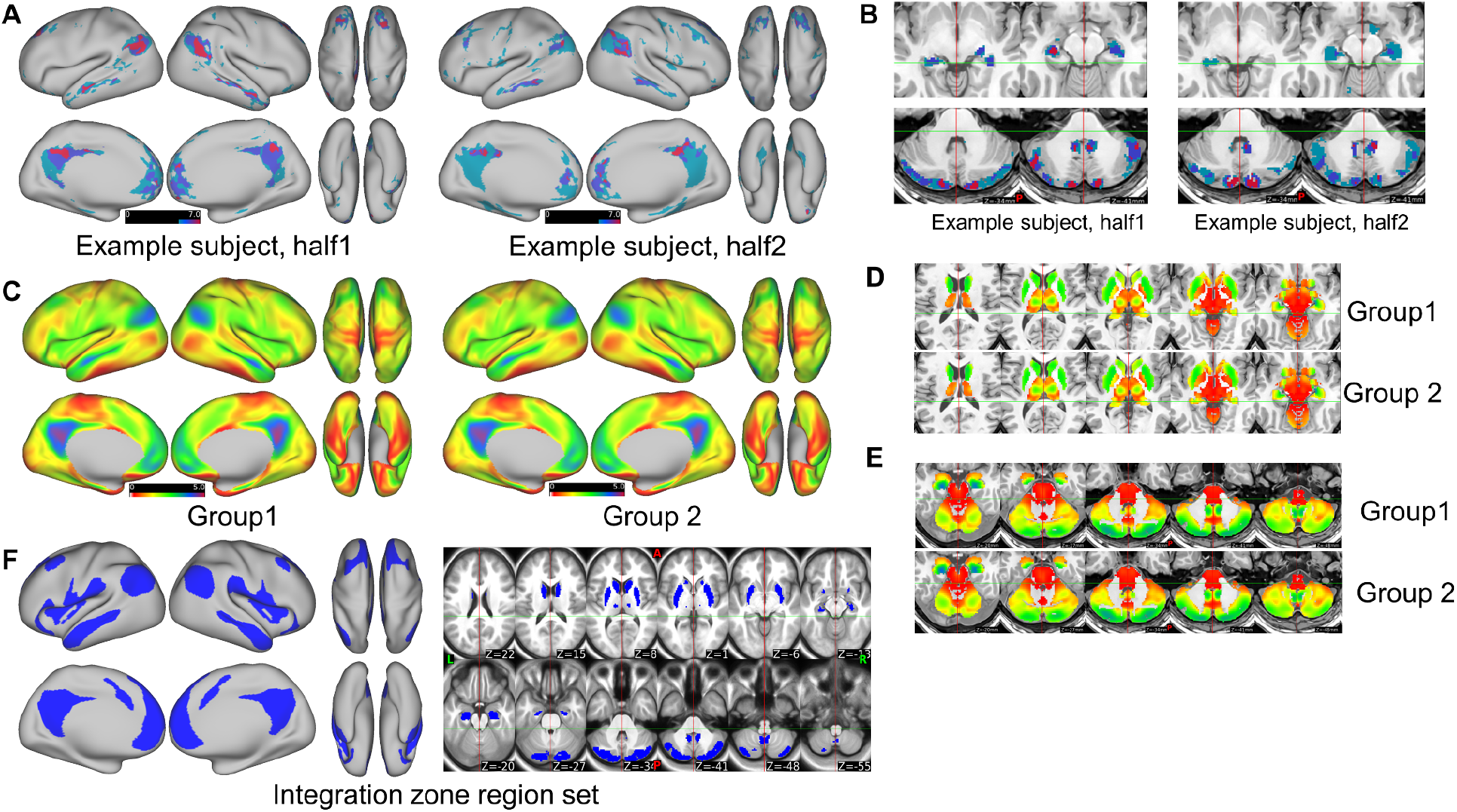
Identifying regions with multiple overlapping networks. A) An example of regions identified on the cortex, subcortical nuclei and **(B)** cerebellum that have 5 or more networks overlapping in an individual subject (image has been thresholded for visualization purposes). C-E) The number of networks that overlap at each grayordinate for groups 1 and 2. The hippocampi, the posterior cerebellum, (in particular the spinocerebellum), also demonstrate high network overlap. **F)** Brain-wide maps the average number of overlapping networks for ABCD group1 (shown in **C**) were thresholded at 2.2 networks to generate an integration zone region set.

Furthermore, integration zones across the population are highly reliable. The number of networks detected at each grayordinates was calculated for ABCD group1 and group2 (see figure 7C). We found that the integration zones across the population were highly reliable (r(91282)=0.9994, p <0.001. We observed that regions with the highest number of networks closely resembled the default mode network (Figure 7), including regions such as the parieto-occipital junction, middle temporal gyrus, posterior cingulate cortex/precuneus, hippocampus, and the posterior aspect of the posterior cerebellum, consistent with prior work in adults (Buckner et al., 2009).

When we used integration zones to perform an identical subset reliability analysis using subsets of participants, we found that integration zones could provide more reproducible statistical maps of executive function brain-wide associations compared to using either the MIDB Probabilistic parcellation or the Gordon parcellation(Figure 4D). To ensure that the improvement in reproducibility was not due to fewer ROIs in the integration zone parcellation, we conducted an additional analysis subset reliability analysis using the Gordon Parcellation, where we randomly sampled the same number of ROIs as the integration zone parcellation. We found that the rise to maximum of the randomly-sampled Gordon parcellation was nearly identical to the complete Gordon parcellation (Figure 4) (supplementary table 5).

### THE MIDB PRECISION BRAIN ATLAS

The MIDB Precision Brain Atlas includes an online tool (https://midbatlas.io) with publicly available ROI sets based on the probability of the neural networks for various methods described here. ROI sets are generated for integration zones and each network separately in 0.005 probability increments. In addition, ROI sets for the combined networks in one label file at the same probability increments are available.

## DISCUSSION

Investigations into brain function, and in particular developmental brain function, requires confidence in structure- and function-based parcellations that consider the vast heterogeneity in functional topography from person to person. The MIDB Precision Brain Atlas provides an invaluable resource to explore the brain function for basic and clinical research that accounts for this individual variation in network topography.

The inaugural MIDB Precision Brain Atlas data comes from the ABCD dataset, which consists of high fidelity individual-specific functional networks for roughly 6000 9-10 year old children with 10 minutes of low-motion resting-state fc-MRI data, along with their associated probabilistic and integration zone maps. The atlas also contains replicas of these data for approximately 9000 subjects generated with concatenated tasks and rest data. Furthermore, we have created an online repository where users can adjust the probability threshold to customize the ROI sets to the desired network probability. We encourage the community to explore the collection of individual precision maps, probabilistic maps, and integrative zones generated from individual-specific networks to characterize how individual variation may impact traditional efforts in mapping network organization to complex behaviors and general network topology across the population.

### Improving reliability in neuroimaging

Some of the noise present in brain-wide association studies is due to sampling variability and random noise in BOLD fluctuations(Feczko et al., 2021; Marek et al., 2020) . However, not accounting for individual topographies in analyses is likely further contributing to systematic noise in rs-fMRI measurements, leading to reduced effect sizes and power. ROIs derived from probabilistic maps demonstrated superior reliability to those based on group averaging. This increased signal-to-noise ratio (SNR) is likely due, in part, to the omission of voxels or grayordinates that demonstrate high network assignment variability across the population (e.g. dorsolateral prefrontal cortex and temporoparietal junction).

The increased SNR provided by using the probabilistic ROI set allows for additional explanatory reliability when conducting BWAS (Figure 4C-E). Accounting for individual-specific topography improves reliability with smaller sample sizes and has the potential to increase effect sizes for some investigations (*but not all*), therefore saving recruitment of potentially hundreds of fewer subjects and hundreds of thousands of dollars in MRI scanning costs.

One way to leverage precision mapping in individuals to increase reliability for group studies is to create probabilistic network description region sets. Probability maps have been used in the structural literature for years, however, there have been limited efforts to produce probabilistic atlases of functional networks, e.g. (Dworetsky et al., 2020; Gordon et al., 2017b). Others have implemented a group-guided methodology to improve detection of functional networks by component-based analysis (Du and Fan, 2013; Harrison et al., 2015; Li et al., 2017). These methods typically force subject-specific functional networks to have component weights that are similar to the group representations, which can dilute subject-specific differences in topography. Recent non-negative matrix factorization (NMF) methods decompose the timeseries into a set of additive parts-based spatial components, yielding a probabilistic parcellation that can be discretized for each subject based on maximal loading to produce individual-specific networks (Cui et al., 2020; Kong et al., 2019). This approach contrasts with TM because we first generate a correlation matrix, then measure the spatial similarity of each grayordinate’s connectivity to a set of networks identified in a group average. By using an approach that leverages the spatial similarity of known networks, we can potentially capture subject-specific functional networks even if subjects demonstrate atypical connection strengths, which has been observed in children with neurodevelopmental disorders (Faraone et al., 2015; Hermosillo et al., 2020).

### Network-specific probabilistic maps have several pros and cons

Structurally-informed parcellations, such as the Desikan parcellation (Desikan et al., 2006; Klein and Tourville, 2012), Destrieux parcellation (Destrieux et al., 2010), M-CRIB (Alexander et al., 2017, 2019), and the Human connectome project (HCP) atlas (Glasser et al., 2016), may not reflect underlying functional network topography in all cases (Figure 4B). One major limitation of these parcellations, as well as the Gordon parcellation (333 parcels within 10 networks) (Gordon et al., 2016), is the assumption that a given parcel participates in the same network in all individuals (Figure 4A). Individual-specific topography confounds this assumption about network assignments. Moreover, atlases that impose network assignments based on gyral-based neuroanatomy likely perpetuate the misconception that identical functions occur at identical locations across individuals, despite obvious intersubject variation in both gyral anatomy and functional connectivity. Analyses that assume identical network assignments across individuals, based on structurally-derived parcellations therefore introduce 2 sources of noise: 1) noise from the misalignment of structural parcellation-to-functional network (Kong et al., 2021) and 2) intersubject network topographic variability. A potential byproduct of including these kinds of noise is that studies require larger sample sizes to observe an effect of a similar magnitude.

By considering individual network topographies and/or focusing on areas that are highly consistent across individuals, one may be able to improve power in large-scale studies by limiting the contribution of individual differences to support inferences about the group. The trade-off in the case of the probabilistic regional mappings is that the sparse brain coverage might obscure important information processing that occurs at these omitted variable locations. In the case of purely predictive analyses, the sparsity of the region set by its nature reduces the feature set able to be included in the prediction; thus, while on the one hand, signals are more reliable relative to traditional region sets, on the other hand a lot of information goes unused, such that even though the “unused regions” are less reliable, they might be important to maximize prediction. Therefore, usage of the MIDB probabilistic ROIs may not be appropriate for all situations.

One important potential usage of probabilistic atlases relates to functional neuronavigation for targeted brain stimulation. Historically, anatomical coordinates, landmarks, or the spatial location of task-based activations in fMRI, have been used to guide non-invasive brain stimulation, such as transcranial magnetic stimulation (TMS). Several lines of work have recently suggested that outcomes might be improved with consideration of personalized functional connectivity (Cole et al., 2021). Thus, Recent advances in brain stimulation using TMS have shifted focus from anatomical brain landmarks to personalized fMRI or functional connectivity with the goal of increasing treatment efficacy (Cash et al., 2020; Fox et al., 2012b). For example, Cash and colleagues found that the variation in the dorsolateral prefrontal cortex (DLPFC) (r)TMS stimulation site location affects antidepressant response (Cash et al., 2019). Specifically, when rTMS stimulation was delivered at sites of the DLPFC per individual that displayed a stronger negative correlation with the subgenual cortex, the antidepressant treatment showed better outcomes (Cash et al., 2019, 2020; Fox et al., 2012b; Weigand et al., 2018). However, as noted throughout, what would a clinician or investigator do to maximize stimulation without sufficient data to generate an individual map, or if no MRI machine is available? Probabilistic mapping provides a potential use case to maximize targeting across a population. Similar to findings from Cash and colleagues (Cash et al., 2019, 2020, 2021a), we demonstrate that a seed placed within a region of high network probability (0.75 probability of frontoparietal) within the DLPFC showed consistent anticorrelation with the subgenual cortex, both in the MSC and ABCD subjects (Figure 8A). However, when the seed was moved slightly outside of the region of high network consensus to a region with high network heterogeneity (0.35 probability), the correlation with the subgenual cortex was inconsistent(Figure 8B). This suggests that the MIDB Probabilistic parcellation allows investigators to quantify the confidence of the spatial location of a network of interest and refine targets for therapeutic brain stimulation, in situations where personalized network maps are not available.

**Figure 8.**
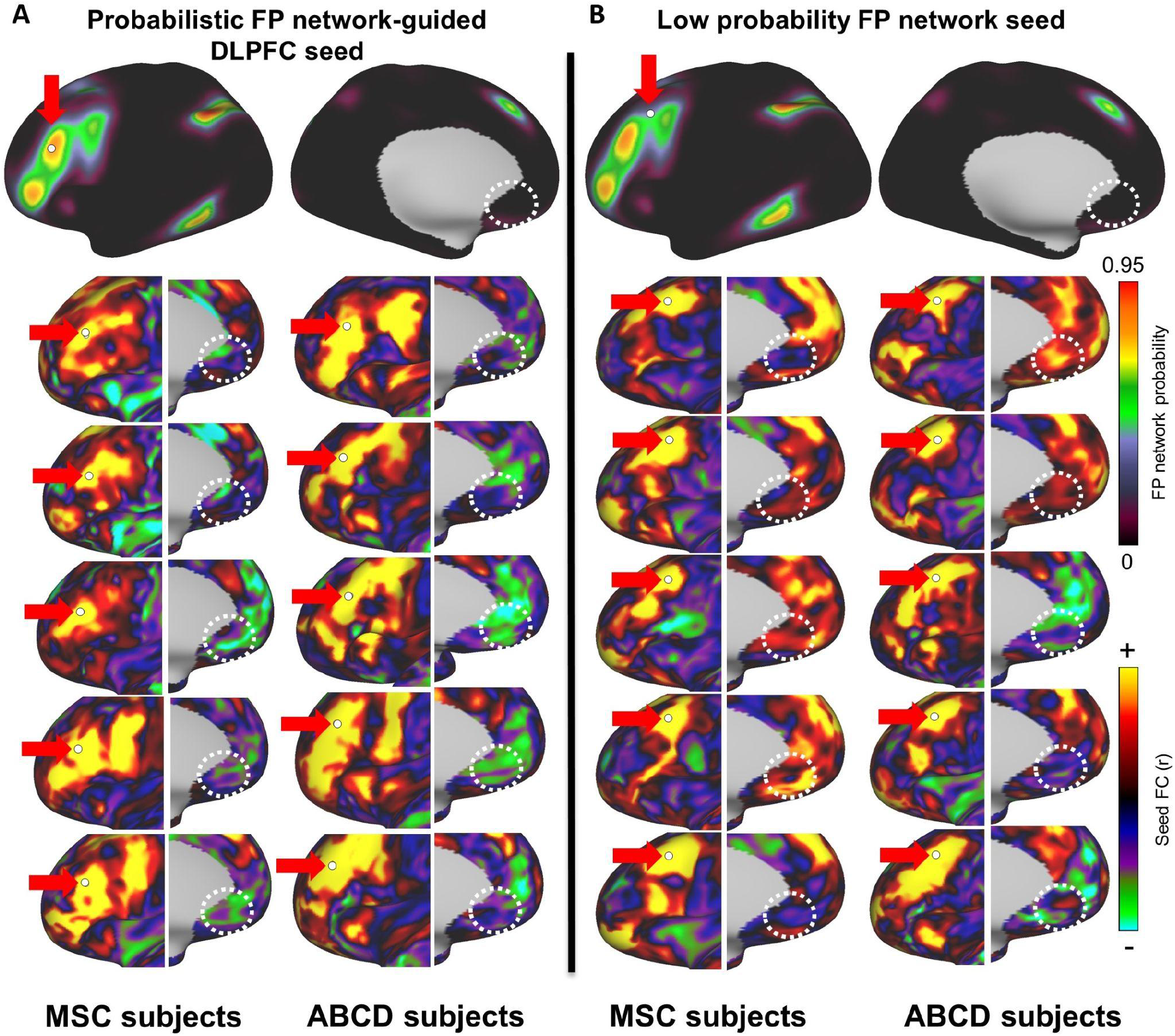
Probabilistic map-guided seed-based correlation. A seed based correlation was conducted with 5 MSC and 5 ABCD subjects. **A)** A seed was placed at the DLPFC (as defined by the MIDB Frontoparietal probabilistic map) and connectivity to the subgenual cortex was examined (dotted white circle). Note that, in most subjects, the connectivity to the frontoparietal network was anti-correlated with the subgenal cortex (green-blue). **B)** When the seed was placed in a region with a low probability of belonging to the frontoparietal network, connectivity to the subgenual cortex was inconsistent. White circles indicate the location of the subgenual gyrus (subgenal MNI coordinates=±5, 25, -10 (Kelly et al., 2009; Zhou et al., 2016)).

### Integregration zones show hub like properties

The MIDB Precision Brain Atlas also includes integration zones (IZ) that represent overlapping networks as determined by the overlapping TM method. We posit that these integration zones are functionally similar to network hubs (Power et al., 2013a; Sporns et al., 2007) (i.e. nodes that have a higher degree of connectedness) and likely play a crucial role in relaying information brain networks. Recent work by Bagarinao and colleagues (Bagarinao et al., 2020) quantified a functional connectivity overlap ratio (FCOR) to examine the spatial extent to which each region belongs to a given network. Regions belonging to several networks (e.g. posterior parietal and posterior cingulate) closely match those that we identified (Figure 7)(Buckner et al., 2009). Due to their central role in fundamental cognitive processes such as attention and consciousness (Silasi and Murphy, 2014), the core features of these integration zones are likely shared across the population, and provide strong between-group reliability (Figure 4D-F). This ROI set can be used to examine the mechanisms of information integration and relay, and targeted brain stimulation (Lynch et al., 2019).

Others have also examined connectivity between integration zones, or more broadly as “hubs” (Gratton et al., 2018b; van den Heuvel and Sporns, 2013; Power et al., 2013a). Hubs that have been previously shown very closely align with the integration zones that we’ve identified here (Buckner et al., 2009). Conventionally, connector hubs have been conceptualized as a category of specific brain regions that allow integration across networks, derived from group averages (Bertolero et al., 2015; Power et al., 2013a). However it is important to note that despite the spatial variability across participants with respect to the number of overlapping brain networks, the location of integration zones was replicated in an independent group. This suggests the appearance of interconnected regions in group data is not an artifact induced by group averaging (Gordon et al., 2018), but is indeed a common feature of network interaction among the population. Furthermore, when we compared the BWAS statistical maps between the randomly sampled Gordon parcellation to the integration zones, we still found a pronounced increase in reproducibility using integration zones, which supports the hypothesis that these regions and their interactions, similar to the ‘rich club’ areas (van den Heuvel and Sporns, 2011; Sporns, 2010), may play a role in the instantiation of complex behaviors . It should be noted that the ability to detect grayordinates that overlap with regard to network assignments, may be obscured by limited resolution (here 2.4 mm isovoxel). Such resolution by nature blurs independent neuronal signals (Supplementary Figure S12) and might artificially lead to overlapping networks or hubs(Braga et al., 2019). Nevertheless, regions with a high density of networks appear to be consistent in the population. Furthermore, neurons that reside at the internetwork boundary likely maintain the boundary through persistent internetwork communication (Carmichael and Price, 1996). See Supplementary text for a discussion of “Volumetric averaging and integration zones”. Thus, integration zones, while still requiring investigations of their origin, are likely important for the integration of information processing across systems.

### Network maps are only one snapshot of topography during development

The ABCD Study data set provides a unique opportunity to explore neural networks longitudinally in a set of racially and ethnically diverse young participants, closely representative of the U.S population (Casey et al., 2018). To our knowledge, this is the first study that has attempted to quantify network topography for a sample of this magnitude with the potential to follow the participants into young adulthood. While we do not anticipate large changes in network topographies from adolescence to adulthood, there is some suggestion that refinements around borders can occur (Cui et al., 2020). The ability to quantify topographies provides a rich avenue for investigation to capture subtle changes over time when compared with metrics such as substance abuse, mental health (Goldstone et al., 2020; Janiri et al., 2020; Karcher et al., 2019; Pagliaccio et al., 2020), neurocognition (Marek et al., 2019), development, and environment (Guerrero et al., 2019; Marshall et al., 2020) in the same cohort. As participants age, the MIDB Precision Brain Atlas will provide age-specific maps.

### MIDB Open Science Framework

The MIDB Precision Brain Atlas is an evolving resource, and we invite the scientific community to contribute toward the additional characterization of brain maps. To start, we are providing up to a dozen network brain maps per participant from the ABCD dataset using various methods of brain mapping, combinations of minutes used (e.g. rest vs task and rest combined), and methodologies. The MIDB Precision Brain Atlas currently includes functional network maps from the ABCD year 1 dataset (Feczko et al., 2021), the MSC dataset (Dworetsky et al., 2020; Gordon et al., 2017a), the HCP dataset (Dworetsky et al., 2020; Van Essen et al., 2012), the Yale Low-res data set (Dworetsky et al., 2020; Scheinost et al., 2016), and the Dartmouth Gordon parcellation dataset (Dworetsky et al., 2020; Gordon et al., 2016). As new techniques for individual-specific brain mapping are developed and integrated into the MIDB Precision Brain Atlas, this will contribute to the development of a highly comprehensive brain mapping resource.

The MIDB Precision Brain Atlases will be an evolving repository of processing and analysis tools and parcellations that are overseen by community partners. All individual-specific maps for ABCD will be downloadable through the National Data Archive (NDA) https://nda.nih.gov/. All others will be downloadable through the website (per each dataset’s usage agreement). For this, investigators who wish to share individual-specific maps based on ABCD data, can do so via the ABCD-BIDS Community Collection (ABCCC; NDA Collection 3165)(Feczko et al., 2020, 2021). Review and inclusion of data into the MIDB Precision Brain Atlas will utilize the community governance structure (https://bids.neuroimaging.io/governance.html). Briefly, the data must be: 1) published with a clear description of the open-source and reproducible tools used to analyze the data, 2) BIDS-formatted, and 3) de-identified so as not to risk reidentification of any participants. Those interested in contributing their code as an ABCD utility can link their repository to the ABCD open science framework (Feczko et al., 2020) after receiving approval. Additional probabilistic maps generated from HCP, MSC, and others are already included in our online tool (Dworetsky et al., 2020).

We hope that the thousands of network maps based on multiple validated methodologies, and replicable population-level probabilistic topographies we are providing will serve as a new avenue of investigation into adolescent development. Furthermore, the high reliability observed from integration zones, merits further investigation as a explanatory source of behavior. As a community atlas, the MIDB Precision Brain Atlas enables the systematic investigation of the contributions of network topography and network-network interaction to human cognition and behavior.

## ACKNOWLEDGEMENTS

This research was supported by National Institutes of Health (NIH) of the United States, National Institute of Mental Health (NIMH) grants awarded to (MH096773 (Fair), MH115357 (Fair)) and a diversity supplement (R01MH115357-02S1 (Hermosillo), R01MH120482 (Satterthwaite), R01EB022573 (Satterthwaite), 37MH125829 (Satterthwaite)). This project made use of Connectome DB and Connectome Workbench, developed under the auspices of the Human Connectome Project at Washington University in St. Louis and associated consortium institutions (http://www.humanconnectome.org/).

The ABCD Study, supported by the National Institutes of Health, is a multisite, longitudinal study designed to recruit more than 10,000 children aged 9-10 years and follow them over 10 years into early adulthood. MRI and Demographic data used in this article were collected by the Adolescent Brain Cognitive Development (ABCD) Study (https://abcdstudy.org), and retrieved from in the NIMH Data Archive (NDA). Data used in the preparation of this article were obtained from the Adolescent Brain Cognitive DevelopmentSM (ABCD) Study (https://abcdstudy.org), held in the NIMH Data Archive (NDA). The ABCD Study® is supported by the National Institutes of Health and additional federal partners under award numbers U01DA041048, U01DA050989, U01DA051016, U01DA041022, U01DA051018, U01DA051037, U01DA050987, U01DA041174, U01DA041106, U01DA041117, U01DA041028, U01DA041134, U01DA050988, U01DA051039, U01DA041156, U01DA041025, U01DA041120, U01DA051038, U01DA041148, U01DA041093, U01DA041089, U24DA041123, U24DA041147. A full list of supporters is available at https://abcdstudy.org/federal-partners.html. A listing of participating sites and a complete listing of the study investigators can be found at https://abcdstudy.org/consortium_members/. ABCD consortium investigators designed and implemented the study and/or provided data but did not necessarily participate in the analysis or writing of this report. This manuscript reflects the views of the authors and may not reflect the opinions or views of the NIH or ABCD consortium investigators.

## STAR METHODS

| Reagent or Resource | Source | Identifier |
| --- | --- | --- |
| Available data | NDA | https://github.com/ABCD-STUDY/nda-abcd-collection-3165 |
| Custom template matching code | This paper | https://gitlab.com/Fair_lab/compare_matrices_to_assign_networks |
| MATLAB | Mathworks | https://www.mathworks.com/ |
| Connectome Workbench | Marcus et al. 2011(Marcus et al., 2011) | http://www.humanconnectome.org/software/connectome-workbench |
| Freesurfer | (Fischl, 2012) | https://surfer.nmr.mgh.harvard.edu/ |
| Infomap | Rosvall and Bergstrom, 2008 (Rosvall & Bergstrom, 2008) | www.mapequation.org |
| FSL | (Smith et al., 2004) | https://fsl.fmrib.ox.ac.uk/fsl/fslwiki |
| Midnight Scan club | (Gordon, Laumann, Gilmore, et al., 2017) | https://www.openfmri.org/dataset/ds000224/ |
| MiDB ROI tool | This paper | https://midbatlas.io/ |
| Subject Brain maps for ABCD collection | This paper | https://nda.nih.gov/ |
| ABCC Collectionfor matched groups | Feczko et al. 2021 | https://www.biorxiv.org/content/10.1101/2021.07.09.451638v1 |

## METHODS

### Participant information

Participants were recruited under the auspice of the ABCD study to follow brain development and health in a longitudinal manner from 9-10 years of age to adolescence.

### MRI image acquisition

MRI images were collected across 21 sites across the United States of America (Children’s Hospital Los Angeles, University of Colorado Boulder, Florida International University, Laureate Institute for Brain Research, Medical University of South Carolina, Oregon Health and Science University, University of Rochester, SRI International, University of California Los Angeles, University of California San Diego, University of Florida, University of Maryland at Baltimore, University of Michigan, University of Minnesota, University of Pittsburgh Medical Center, University of Utah, University of Vermont, University of Wisconsin-Madison, Virginia Commonwealth University, and Washington University in St. Louis) (Casey et al., 2018). The imaging component of the study was developed by the ABCD Data Analysis and Informatics Center (DAIC) and the ABCD Imaging Acquisition Workgroup.

Sequences were harmonized across Siemens, Philips, and GE 3T scanners. For further detail regarding MRI acquisitions, see (Casey et al., 2018). Briefly, subjects underwent 25-45 minutes of pre-scan task compliance, localizer, 3d T1-weighted MRI (1mm isotropic, TR=(either 2500 or 6100 ms, TE=2-2.9 ms ,8° flip angle, 256 x 256 FOV), diffusion weighted images, 3d T2-weighted MRI (1mm isotropic, TR=2500 or 3200ms, TE=60–565ms, variable flip angle, 256 x 256 FOV), 1-2 runs of rs-fMRI (1mm isotropic, TR=800ms, TE=30, variable flip angle= 52°, 216 x 216 FOV), and a randomized order of monetary incentive delay (MID), stop signal task (SST), and emotional n-back (EN-back) tasks.

Of the original 11572 participants from the ABCD 2.0 release(Volkow et al., 2018), participants were divided in to a discovery (n=5786) and replication (n=5786) set that were matched along 10 variables: site location, age, sex, ethnicity, grade, highest level of parental education, handedness, combined family income, and exposure to anesthesia (Marek et al., 2019) (see Supplementary Table 1). All resting state scans were acquired using a gradient-echo EPI sequence (TR =800 ms, TE =30 ms, flip angle = 90°, voxel size = 2.4 mm3, 60 slices). Head motion was monitored in real time using Framewise Integrated Real-time MRI Monitor (FIRMM) software at Siemens sites (Dosenbach et al., 2017). For resting state scans, participants were instructed to lie still and fixate on a crosshair at the center of their visual field.

All functional MRIs were processed with the publicly available ABCD-BIDS pipeline (https://github.com/DCAN-Labs/abcd-hcp-pipelines), which is a modified version of the HCP processing pipelines(Feczko et al., 2021). Brain extraction was performed by PreFreesurfer after denoising and bias field correction of the anatomical T1 and/or T2 weighted images. The DCAN-labs processing pipeline applies ANTs DenoiseImage to improve structural clarity and ANTs N4BiasFeildCorrection (Advanced Normalization Tools) (Avants et al., 2009; Tustison et al., 2010) to reduce field bias (Marek et al., 2019).

### Resting state time course processing

#### Signal regression

Time courses were corrected using DCAN-BOLDproc (Feczko et al., 2021). The method for signal regression has been previously described (Hermosillo et al., 2020). Briefly, resting state time courses (using surface registration for cortex and volume registration for subcortical gray matter) were detrended and further processed using mean whole brain, ventricle, and white matter signal as well as displacement on the 6 degrees of freedom, rigid body registration, their derivatives and their squares by regression (Ciric et al., 2017; Friston et al., 2000; Power et al., 2014). Lastly, time courses were filtered using a first order Butterworth band pass filter between 9 and 80 mHz backwards and forwards using MATLAB’s *filtfilt* function (v2016-2018x The MathWorks, Cambridge, UK).

The BOLD fMRI volumetric data from the cerebral cortex was constrained to the cortical sheet for surface-based imaging (Glasser et al., 2013) and combined with volumetric midbrain and hindbrain time courses into a CIFTI format. Once BOLD data was mapped to the sheet, time courses were deformed and resampled to the original surface.

#### Head motion correction

Head movement in the scanner interferes with the ability to identify a grayordinate from one time point to the next and the movement of a large electrically conductive tissue in a magnet introduces contaminating artifacts from eddy currents. To minimize these effects, we rigorously controlled for head motion by using a framewise displacement threshold of 0.2 mm and only using subjects with at least 10 minutes of resting state data post-motion correction. Movement was calculated by framewise displacement (FD) in mm) using the formula *FD_i_* = ∣Δ*d_ix_*∣ + ∣Δ*d_iy_*∣ + ∣Δ*d_iz_*∣ + ∣Δα*_i_*∣ + ∣Δβ*_i_*∣ + ∣Δ_γ*i*_∣, where Δ*d_ix_* is the frame-to-frame change in the x position: Δ*d_ix_* = *d*_(*i*−1)*x*_ − *d_ix_*, and so forth for the other rigid body parameters [*d_ix_*, *d_iy_*, *d_iz_*, α*_i_*, β*_i_*, γ*_i_*] (Power et al., 2013b). Rotational displacements were converted from degrees to millimeters by calculating displacement on the surface of a sphere with a 50 mm radius, which is approximately the mean distance from the cerebral cortex to the center of the head. Frames were removed if their total relative movement in any direction (FD) was greater than 0.2 mm relative to the previous frame or if they were contained within a segment of 5 contiguous frames that violated the threshold.

For the remaining frames, the standard deviation was calculated across all grayordinates to remove potential artifacts. Frames that had outliers in the standard deviation of the bold signal were removed using the Median absolute deviation method in MATLAB and Statistics Toolbox Release 2016b (The Mathworks In., Natick, Massachusetts, United States). In time courses containing more than 10 minutes of resting state data, frames were randomly sampled to generate correlation matrices using exactly 10 minutes of fs-MRI data. Of the 11572 participants enrolled, 10,038 had usable structural and functional MRI collected, and of these based on our movement/signal criteria, approximately 3973 (∼40%) children were excluded based on excessive movement in the scanner during resting state scans. During task-based scans, we were able to retain many more usable frames at the FD criteria (group1: n= 4699, group2: n=4732), excluding only 607 participants (6%).

### Infomap community detection method

The community detection method using the graph theory-based algorithm Infomap has been previously described (Gordon et al., 2017a; Power et al., 2011). The same correlation matrices that were used in the TM processes were used to detect networks using Infomap. Briefly, vertices/voxels within 30 mm of each other were set to zero in the matrix to avoid biasing network membership for nearby connections that had undergone spatial smoothing. The resulting correlation matrix was then thresholded at a range of density thresholds (0.3%, 0.4%, 0.5%, 1%, 1.5%, 2.0%, 2.5%, 3.0%) and each one was used as an input for Infomap. For instances where Infomap was implemented on combined cortical and subcortical data (data shown in Figure 3C and the average group matrix shown in Figure S6), we extended the range of density thresholds to include 4% and 5%. Infomap calculates the network assignment based on an optimized code length using a flow-based method (Rosvall and Bergstrom, 2007, 2008). Networks that are computed in the group average are labeled based on similar patterns of activation observed in the scientific literature (Dworetsky et al., 2020; Gordon et al., 2017a, 2017b). Small networks with 400 or fewer grayordinates were defined as “unassigned”.

Networks identified in each individual were then labelled based on the Jaccard Similarity to a network observed in the group average, however, often individuals will retain novel networks that are not observed in group averaging, and these remained unlabeled. The list of networks included are the default mode network (DMN), the visual network (VIS), the frontal parietal network (FPN), The premotor network (PMN), the dorsal attention network (DAN), the ventral attention network (VAN), the salience network (Sal), the cingulo-opercular network (CO), the sensorimotor dorsal network (SMd), the sensorimotor lateral Network (SMl), the auditory network (AUD), the anterior medial temporal network (AMTL), the posterior medial temporal network (post MTL), parieto-occipital network PON, and the parietal medial network (PMN) (Gordon et al., 2017a). In each subject and in the average, a “consensus” network assignment was determined across the various thresholds, by giving each node the assignment it had at the sparsest possible threshold at which it was successfully assigned to one of the known group networks. Contiguous network clusters that were smaller than 30 grayordinates were removed and merged into neighboring networks, with the largest networks given priority.

### Template matching method

Multiple versions of the time series were used depending on the analysis: either exactly 10 minutes of randomly sampled frames, all available frames below the FD threshold, or concatenated rest and task data in the following order: rest, MID, n-back, and SST (provided that the participant had an available scan for the task). To generate the templates, Infomap community detection was performed at several tie densities (for full details of average networks, see (Gordon et al., 2017a, 2017b; Laumann et al., 2015) on an average connectivity matrix (n=120 participants) using a two level solution. This yielded 14 networks which include: the default mode network (DMN), the visual network (VIS), the frontal parietal network (FPN), the dorsal attention network (DAN), the ventral attention network (VAN), the salience network (Sal), the cingulo-opercular network (CO), the sensorimotor dorsal network (SMd), the sensorimotor lateral network (SMl), the auditory network (AUD), the temporal pole network (Tpole), the medial temporal network (MTL), the parietal occipital network (PON), and the parietal medial network (PMN). Sensory and motor systems were combined due to the coupled nature of activation. Despite high reproducibility in resting state functional connectivity, the extent to which these networks are activated on a neuronal time scale is unclear. However, recent work by Gratton and colleagues suggests that the contribution of short-term dynamic changes (e.g. from task-based states) to variation in brain organization is quite modest relative to resting state organization (Gratton et al., 2018b).

To generate an independent template, we conducted a seed-based correlation (using an average time series correlated to all the grayordinates) for all networks. Seed-based correlations were generated using the dense time series from each template participant that were smoothed with a within-frame spatial Gaussian smoothing kernal of 2.55 mm using each participant’s own midthickness surfaces (extracted from the Surf stage of Freesurfer). The resulting networks were converted to a dlabel CIFTI file and applied to the smooth dense series to generate an average time series for each network. We then correlated the time series of the seed with the times series of all other grayordinates. The seed and remaining time series were motion censored using an FD of 0.2 millimeters and outliers in the BOLD signal were removed using the median absolute deviation in the remaining frames using the motion censoring method outlined above.

Seed-based correlation values were averaged across all the participants in the template group (n=164, 9-10 year olds), resulting in a vector (91282 x 1) of average correlation values for each network correlated with each grayordinate. Each network vector was averaged independently across subjects in the template group to generate seed-based templates for each network. We then thresholded each network template at Z ≥ 1.

To generate precision maps for each participant in ABCD groups 1 and 2, we examined the whole-brain connectivity for each grayordinate by correlating the dense time series against all other grayordinates. For each participant in each test group (group 1 n=∼ 5000, group 2 n=∼ 5000), we generated a Pearson correlation matrix (91282 x 91282 grayordinates) for each connection using the dense time series using the Connectome workbench command “-cifti-correlate” (https://www.humanconnectome.org/software/connectome-workbench). Time series were then motion censored (see motion censoring and supplementary methods) to reduce artifacts induced by head motion.

Because connectivity matrices were generated including subcortical brain regions, the correlation matrix was Z-scored separately for each hemisphere, the subcortical region, and the connections between the cortex and the subcortex. This allowed for normalization of connectivity between subcortex and cortex where there is the potential for a decreased signal-to-noise ratio in the subcortex. We threshold the whole-brain connectivity for each grayordinate to only include correlated grayordinates with Z-scores values greater than or equal to one. This resulted in a vector of whole-brain connectivity for each grayordinate that only includes grayordinates that are strongly correlated to a given network template. We then calculate an eta^2^ value between the remaining grayordinates and each of the network templates seen in Supplementary Figure S4. The grayordinate is assigned to whichever network with the maximum eta^2^ value.

### Overlapping template matching method

To generate overlapping networks for each participant, rather than assigning the grayordinate to the network with the maximum eta^2^ value, we used a data-driven approach to assign multiple networks to each grayordinate. For each network we plotted the distribution of eta^2^ values (Figure 5C). The connectivity for each network demonstrates a characteristic skewed bimodal distribution. The distribution for eta^2^ values was distributed into 10,000 bins and fitted with a cubic spline. The distribution was then smoothed using a Savitzky-Golay filter using a 2,000 data point window within MATLAB (v2016-2018x The MathWorks). We calculated the local minimum of the bimodal distribution by taking the derivative of the smoothed data between 4,000 and 7,000 bins. We then used this local minimum as the threshold for whether or not a grayordinate would be labelled with this network, where grayordinates above this threshold would receive the network assignment. Grayordinates that had an eta^2^ value higher than the threshold were assigned to those networks (Figure 5D).

### Probabilistic maps

Probabilistic maps were generated separately for each group, method, and network separately. Probabilistic maps were generated by calculating the probability that a grayordinate was assigned to a given network using all the participants within the group. The TM ROI set was generated by converting clusters produced by thresholding the probability maps at 0.8, (excluding clusters smaller than 30 grayordinates), converting them to dlabel files (dlabel.nii), and combining ROIs into 1 combined probabilistic parcellation. Probabilistic dlabel files are available for the combined networks and each network separately from the MiDB Precision Brain Atlas webpage: http://neuroatlas.org.

### NMF community detection method

We implemented a community detection technique used previously to decompose non-negative subject-specific functional networks using their corresponding concatenated rest+task dense time series in a constrained manner using 3 regularized terms (Cui et al., 2020; Lee and Seung, 1999; Li et al., 2017). Briefly, a voxel-wise group sparsity regularization term was first used to ensure a group consensus was used as a prior using Group 3. Second, spatial locality regularization term was used to ensure that functional coherent voxels are encouraged to reside in the same functional network. Lastly, a within-subjects regularization term was used to eliminate redundant functional networks (Cui et al., 2020; Li et al., 2017). The weights from the consensus were then applied to each of the time series for subjects in ABCD groups 1 and 2.

### Analysis of minutes necessary for reliable communities using split halves

We calculated the similarity between split halves for an individual by splitting the resting state time series in half and generating a correlation matrix of all grayordinates from each half using exactly 10 minutes of randomly sampled frames. Each correlation matrix was used as an input to both the TM algorithm and Infomap algorithm to generate networks for each half (see Methods: Template matching technique). We then calculated the normalized mutual information (NMI) between halves (https://github.com/MidnightScanClub/MSCcodebase). To create the null distribution we calculated the NMI between an individual subject’s half and all other halves in the group set for all subjects. The difference between the test (self) and null (other) distributions was assessed using an independent two-sample t-test with unequal variance.

### Brain-Behavior associations using subset reliability

To assess the reliability of a probabilistic parcellation schema, we conducted a split-group subset reliability association analysis. We randomly sampled participants from Group 1 at discrete sample sizes and correlated each corresponding element of the matrix to measure reliability against the subjects’ behavioral measures. For each analysis we quantified the correlation between each subject’s behavioral measure and 1) Gordon connectivity matrix, 2) probabilistic parcellation connectivity matrix, or 3) integrative zone. The resultant correlation matrix for each subset was then correlated to the correlation matrix made from all the participants in Group 2. To get nonlinear regression estimate across sample sizes, we then fitted a curve through the data points using an Exponential Rise to Maximum Single 3-Parameter estimate with the following equation:

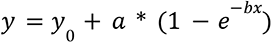

where *y* is the correlation, *y*_0_ is the y-intercept, *a* is a scaling parameter, *b* is the rate of rise to maximum and x is the number of subjects. SigmaPlot 12.5 (Systat Software, San Jose, CA). All regression parameter fits were significant (p<0.0001) and were highly correlated with the data (PC1 Gordon: r^2^=0.8045, TM: 0.8540; PC2: Gordon=0.7181, TM=0.6260; PC3: Gordon=0.8086, TM=0.6999).

To ensure that the increase in intergroup reproducibility observed with the integration zones was not simply due to the reduced number of ROIs, we conducted an additional subset reliability analysis, where we randomly sampled 30 ROIs (the same number of ROIs in the integration zone set) at each of the various subject subsets (Figure 4D-F).

## REFERENCES

Alexander, B., Murray, A.L., Loh, W.Y., Matthews, L.G., Adamson, C., Beare, R., Chen, J., Kelly, C.E., Rees, S., Warfield, S.K., et al. (2017). A new neonatal cortical and subcortical brain atlas: the Melbourne Children’s Regional Infant Brain (M-CRIB) atlas. Neuroimage 147, 841–851.

Alexander, B., Loh, W.Y., Matthews, L.G., Murray, A.L., Adamson, C., Beare, R., Chen, J., Kelly, C.E., Anderson, P.J., Doyle, L.W., et al. (2019). Desikan-Killiany-Tourville Atlas Compatible Version of M-CRIB Neonatal Parcellated Whole Brain Atlas: The M-CRIB 2.0. Front. Neurosci. 13, 34.

Andersen, R.A. (1997). Multimodal integration for the representation of space in the posterior parietal cortex. Philos. Trans. R. Soc. Lond. B Biol. Sci. 352, 1421–1428.

Avants, B.B., Tustison, N., and Song, G. (2009). Advanced normalization tools (ANTS). Insight J. 2, 1–35.

Bagarinao, E., Watanabe, H., Maesawa, S., Mori, D., Hara, K., Kawabata, K., Ohdake, R., Masuda, M., Ogura, A., Kato, T., et al. (2020). Identifying the brain’s connector hubs at the voxel level using functional connectivity overlap ratio. Neuroimage 222, 117241.

Bertolero, M.A., Yeo, B.T.T., and D’Esposito, M. (2015). The modular and integrative functional architecture of the human brain. Proc. Natl. Acad. Sci. U. S. A. 112, E6798–E6807.

Braga, R.M., and Buckner, R.L. (2017). Parallel Interdigitated Distributed Networks within the Individual Estimated by Intrinsic Functional Connectivity. Neuron 95, 457–471.e5.

Braga, R.M., Van Dijk, K.R.A., Polimeni, J.R., Eldaief, M.C., and Buckner, R.L. (2019). Parallel distributed networks resolved at high resolution reveal close juxtaposition of distinct regions. J. Neurophysiol. 121, 1513–1534.

Brodmann, K. (1909). Vergleichende lokalisationslehre der grosshirnrinde in ihren prinzipien dargestellt auf grund des zellenbaues (J.A. Barth).

Buckner, R.L., Sepulcre, J., Talukdar, T., Krienen, F.M., Liu, H., Hedden, T., Andrews-Hanna, J.R., Sperling, R.A., and Johnson, K.A. (2009). Cortical Hubs Revealed by Intrinsic Functional Connectivity: Mapping, Assessment of Stability, and Relation to Alzheimer’s Disease. Journal of Neuroscience 29, 1860–1873.

Carmichael, S.T., and Price, J.L. (1996). Connectional networks within the orbital and medial prefrontal cortex of macaque monkeys. J. Comp. Neurol. 371, 179–207.

Casey, B.J., Cannonier, T., Conley, M.I., Cohen, A.O., Barch, D.M., Heitzeg, M.M., Soules, M.E., Teslovich, T., Dellarco, D.V., Garavan, H., et al. (2018). The Adolescent Brain Cognitive Development (ABCD) study: Imaging acquisition across 21 sites. Dev. Cogn. Neurosci. 32, 43–54.

Cash, R.F.H., Zalesky, A., Thomson, R.H., Tian, Y., Cocchi, L., and Fitzgerald, P.B. (2019). Subgenual functional connectivity predicts antidepressant treatment response to transcranial magnetic stimulation: independent validation and evaluation of personalization. Biol. Psychiatry 86, e5–e7.

Cash, R.F.H., Weigand, A., Zalesky, A., Siddiqi, S.H., Downar, J., Fitzgerald, P.B., and Fox, M.D. (2020). Using brain imaging to improve spatial targeting of TMS for depression. Biol. Psychiatry.

Cash, R.F.H., Cocchi, L., Lv, J., Fitzgerald, P.B., and Zalesky, A. (2021a). Functional Magnetic Resonance Imaging–Guided Personalization of Transcranial Magnetic Stimulation Treatment for Depression. JAMA Psychiatry 78, 337–339.

Cash, R.F.H., Cocchi, L., Lv, J., Wu, Y., Fitzgerald, P.B., and Zalesky, A. (2021b). Personalized connectivity-guided DLPFC-TMS for depression: Advancing computational feasibility, precision and reproducibility. Hum. Brain Mapp.

Churchland, P.S., and Sejnowski, T.J. (1988). Perspectives on cognitive neuroscience. Science 242, 741–745.

Ciric, R., Wolf, D.H., Power, J.D., Roalf, D.R., Baum, G.L., Ruparel, K., Shinohara, R.T., Elliott, M.A., Eickhoff, S.B., Davatzikos, C., et al. (2017). Benchmarking of participant-level confound regression strategies for the control of motion artifact in studies of functional connectivity. Neuroimage 154, 174–187.

Cole, E.J., Phillips, A.L., Bentzley, B.S., Stimpson, K.H., Nejad, R., Barmak, F., Veerapal, C., Khan, N., Cherian, K., Felber, E., et al. (2021). Stanford Neuromodulation Therapy (SNT): A Double-Blind Randomized Controlled Trial. Am. J. Psychiatry appiajp202120101429.

Cui, Z., Li, H., Xia, C.H., Larsen, B., Adebimpe, A., Baum, G.L., Cieslak, M., Gur, R.E., Gur, R.C., Moore, T.M., et al. (2020). Individual Variation in Functional Topography of Association Networks in Youth. Neuron 106, 340–353.e8.

Desikan, R.S., Ségonne, F., Fischl, B., Quinn, B.T., Dickerson, B.C., Blacker, D., Buckner, R.L., Dale, A.M., Maguire, R.P., Hyman, B.T., et al. (2006). An automated labeling system for subdividing the human cerebral cortex on MRI scans into gyral based regions of interest. Neuroimage 31, 968–980.

Destrieux, C., Fischl, B., Dale, A., and Halgren, E. (2010). Automatic parcellation of human cortical gyri and sulci using standard anatomical nomenclature. Neuroimage 53, 1–15.

Dosenbach, N.U.F., Koller, J.M., Earl, E.A., Miranda-Dominguez, O., Klein, R.L., Van, A.N., Snyder, A.Z., Nagel, B.J., Nigg, J.T., Nguyen, A.L., et al. (2017). Real-time motion analytics during brain MRI improve data quality and reduce costs. Neuroimage 161, 80–93.

Driver, J., and Noesselt, T. (2008). Multisensory interplay reveals crossmodal influences on “sensory-specific” brain regions, neural responses, and judgments. Neuron 57, 11–23.

Du, Y., and Fan, Y. (2013). Group information guided ICA for fMRI data analysis. Neuroimage 69, 157–197.

Dworetsky, A., Seitzman, B.A., Adeyemo, B., Neta, M., Coalson, R.S., Petersen, S.E., and Gratton, C. (2020). Probabilistic mapping of human functional brain networks identifies regions of high group consensus.

von Economo, C.F., and Koskinas, G.N. (1925). Die cytoarchitektonik der hirnrinde des erwachsenen menschen (J. Springer).

Evans, A.C., Collins, D.L., Mills, S.R., Brown, E.D., Kelly, R.L., and Peters, T.M. (1993). 3D statistical neuroanatomical models from 305 MRI volumes. In 1993 IEEE Conference Record Nuclear Science Symposium and Medical Imaging Conference, pp. 1813–1817 vol.3.

Fair, D.A., Cohen, A.L., Power, J.D., Dosenbach, N.U.F., Church, J.A., Miezin, F.M., Schlaggar, B.L., and Petersen, S.E. (2009). Functional Brain Networks Develop from a “Local to Distributed” Organization. PLoS Comput. Biol. 5, e1000381.

Fan, L., Li, H., Zhuo, J., Zhang, Y., Wang, J., Chen, L., Yang, Z., Chu, C., Xie, S., Laird, A.R., et al. (2016). The Human Brainnetome Atlas: A New Brain Atlas Based on Connectional Architecture. Cereb. Cortex 26, 3508–3526.

Faraone, S.V., Asherson, P., Banaschewski, T., Biederman, J., Buitelaar, J.K., Ramos-Quiroga, J.A., Rohde, L.A., Sonuga-Barke, E.J.S., Tannock, R., and Franke, B. (2015). Attention-deficit/hyperactivity disorder. Disease Primers. Nat. Rev. 15020.

Feczko, E., Earl, E., Perrone, A., and Fair, D. (2020). ABCD-BIDS Community Collection (ABCC) (OSF).

Feczko, E., Conan, G., Marek, S., Tervo-Clemmens, B., Cordova, M., Doyle, O., Earl, E., Perrone, A., Sturgeon, D., Klein, R., et al. (2021). Adolescent Brain Cognitive Development (ABCD) Community MRI Collection and Utilities.

Fonov, V., Evans, A.C., Botteron, K., Almli, C.R., McKinstry, R.C., Collins, D.L., and Brain Development Cooperative Group (2011). Unbiased average age-appropriate atlases for pediatric studies. Neuroimage 54, 313–327.

Fox, M.D., Halko, M.A., Eldaief, M.C., and Pascual-Leone, A. (2012a). Measuring and manipulating brain connectivity with resting state functional connectivity magnetic resonance imaging (fcMRI) and transcranial magnetic stimulation (TMS). Neuroimage 62, 2232–2243.

Fox, M.D., Buckner, R.L., White, M.P., Greicius, M.D., and Pascual-Leone, A. (2012b). Efficacy of transcranial magnetic stimulation targets for depression is related to intrinsic functional connectivity with the subgenual cingulate. Biol. Psychiatry 72, 595–603.

Friston, K.J., Mechelli, A., Turner, R., and Price, C.J. (2000). Nonlinear responses in fMRI: the Balloon model, Volterra kernels, and other hemodynamics. Neuroimage 12, 466–477.

Glasser, M.F., and Van Essen, D.C. (2011). Mapping human cortical areas in vivo based on myelin content as revealed by T1- and T2-weighted MRI. J. Neurosci. 31, 11597–11616.

Glasser, M.F., Sotiropoulos, S.N., Wilson, J.A., Coalson, T.S., Fischl, B., Andersson, J.L., Xu, J., Jbabdi, S., Webster, M., Polimeni, J.R., et al. (2013). The minimal preprocessing pipelines for the Human Connectome Project. Neuroimage 80, 105–124.

Glasser, M.F., Coalson, T.S., Robinson, E.C., Hacker, C.D., Harwell, J., Yacoub, E., Ugurbil, K., Andersson, J., Beckmann, C.F., Jenkinson, M., et al. (2016). A multi-modal parcellation of human cerebral cortex. Nature 536, 171–178.

Goldstone, A., Javitz, H.S., Claudatos, S.A., Buysse, D.J., Hasler, B.P., de Zambotti, M., Clark, D.B., Franzen, P.L., Prouty, D.E., Colrain, I.M., et al. (2020). Sleep Disturbance Predicts Depression Symptoms in Early Adolescence: Initial Findings From the Adolescent Brain Cognitive Development Study. J. Adolesc. Health 66, 567–574.

Gordon, E.M., Breeden, A.L., Bean, S.E., and Vaidya, C.J. (2014). Working memory-related changes in functional connectivity persist beyond task disengagement. Hum. Brain Mapp. 35, 1004–1017.

Gordon, E.M., Laumann, T.O., Adeyemo, B., Huckins, J.F., Kelley, W.M., and Petersen, S.E. (2016). Generation and Evaluation of a Cortical Area Parcellation from Resting-State Correlations. Cereb. Cortex 26, 288–303.

Gordon, E.M., Laumann, T.O., Gilmore, A.W., Newbold, D.J., Greene, D.J., Berg, J.J., Ortega, M., Hoyt-Drazen, C., Gratton, C., Sun, H., et al. (2017a). Precision Functional Mapping of Individual Human Brains. Neuron 95, 791–807.e7.

Gordon, E.M., Laumann, T.O., Adeyemo, B., and Petersen, S.E. (2017b). Individual Variability of the System-Level Organization of the Human Brain. Cereb. Cortex 27, 386–399.

Gordon, E.M., Lynch, C.J., Gratton, C., Laumann, T.O., Gilmore, A.W., Greene, D.J., Ortega, M., Nguyen, A.L., Schlaggar, B.L., Petersen, S.E., et al. (2018). Three Distinct Sets of Connector Hubs Integrate Human Brain Function. Cell Rep. 24, 1687–1695.e4.

Gratton, C., Laumann, T.O., Nielsen, A.N., Greene, D.J., Gordon, E.M., Gilmore, A.W., Nelson, S.M., Coalson, R.S., Snyder, A.Z., Schlaggar, B.L., et al. (2018a). Functional Brain Networks Are Dominated by Stable Group and Individual Factors, Not Cognitive or Daily Variation. Neuron 98, 439–452.e5.

Gratton, C., Sun, H., and Petersen, S.E. (2018b). Control networks and hubs. Psychophysiology 55.

Gratton, C., Kraus, B.T., Greene, D.J., Gordon, E.M., Laumann, T.O., Nelson, S.M., Dosenbach, N.U.F., and Petersen, S.E. (2020). Defining Individual-Specific Functional Neuroanatomy for Precision Psychiatry. Biol. Psychiatry 88, 28–39.

Greene, D.J., Marek, S., Gordon, E.M., Siegel, J.S., Gratton, C., Laumann, T.O., Gilmore, A.W., Berg, J.J., Nguyen, A.L., Dierker, D., et al. (2020). Integrative and Network-Specific Connectivity of the Basal Ganglia and Thalamus Defined in Individuals. Neuron 105, 742–758.e6.

Guerrero, M.D., Barnes, J.D., Chaput, J.-P., and Tremblay, M.S. (2019). Screen time and problem behaviors in children: exploring the mediating role of sleep duration. Int. J. Behav. Nutr. Phys. Act. 16, 105.

Harrison, S.J., Woolrich, M.W., Robinson, E.C., Glasser, M.F., Beckmann, C.F., Jenkinson, M., and Smith, S.M. (2015). Large-scale probabilistic functional modes from resting state fMRI. Neuroimage 109, 217–231.

Hermosillo, R.J.M., Mooney, M.A., Fezcko, E., Earl, E., Marr, M., Sturgeon, D., Perrone, A., Dominguez, O.M., Faraone, S.V., Wilmot, B., et al. (2020). Polygenic Risk Score–Derived Subcortical Connectivity Mediates Attention-Deficit/Hyperactivity Disorder Diagnosis. Biological Psychiatry: Cognitive Neuroscience and Neuroimaging 5, 330–341.

van den Heuvel, M.P., and Sporns, O. (2011). Rich-Club Organization of the Human Connectome. J. Neurosci. 31, 15775–15786.

van den Heuvel, M.P., and Sporns, O. (2013). Network hubs in the human brain. Trends Cogn. Sci. 17, 683–696.

van den Heuvel, M.P., and Sporns, O. (2019). A cross-disorder connectome landscape of brain dysconnectivity. Nat. Rev. Neurosci. 20, 435–446.

Huth, A.G., de Heer, W.A., Griffiths, T.L., Theunissen, F.E., and Gallant, J.L. (2016). Natural speech reveals the semantic maps that tile human cerebral cortex. Nature 532, 453–458.

Janiri, D., Doucet, G.E., Pompili, M., Sani, G., Luna, B., Brent, D.A., and Frangou, S. (2020). Risk and protective factors for childhood suicidality: a US population-based study. Lancet Psychiatry 7, 317–326.

Karcher, N.R., O’Brien, K.J., Kandala, S., and Barch, D.M. (2019). Resting-State Functional Connectivity and Psychotic-like Experiences in Childhood: Results From the Adolescent Brain Cognitive Development Study. Biol. Psychiatry 86, 7–15.

Kelly, A.M.C., Di Martino, A., Uddin, L.Q., Shehzad, Z., Gee, D.G., Reiss, P.T., Margulies, D.S., Castellanos, F.X., and Milham, M.P. (2009). Development of anterior cingulate functional connectivity from late childhood to early adulthood. Cereb. Cortex 19, 640–657.

Keuken, M.C., and Forstmann, B.U. (2015). A probabilistic atlas of the basal ganglia using 7 T MRI. Data in Brief 4, 577–582.

Klein, A., and Tourville, J. (2012). 101 labeled brain images and a consistent human cortical labeling protocol. Front. Neurosci. 6, 171.

Kong, R., Li, J., Orban, C., Sabuncu, M.R., Liu, H., Schaefer, A., Sun, N., Zuo, X.-N., Holmes, A.J., Eickhoff, S.B., et al. (2019). Spatial Topography of Individual-Specific Cortical Networks Predicts Human Cognition, Personality, and Emotion. Cereb. Cortex 29, 2533–2551.

Kong, R., Yang, Q., Gordon, E., Xue, A., Yan, X., Orban, C., Zuo, X.-N., Spreng, N., Ge, T., Holmes, A., et al. (2021). Individual-Specific Areal-Level Parcellations Improve Functional Connectivity Prediction of Behavior.

Laumann, T.O., Gordon, E.M., Adeyemo, B., Snyder, A.Z., Joo, S.J., Chen, M.-Y., Gilmore, A.W., McDermott, K.B., Nelson, S.M., Dosenbach, N.U.F., et al. (2015). Functional System and Areal Organization of a Highly Sampled Individual Human Brain. Neuron 87, 657–670.

Lee, D.D., and Seung, H.S. (1999). Learning the parts of objects by non-negative matrix factorization. Nature 401, 788–791.

Li, H., Satterthwaite, T.D., and Fan, Y. (2017). Large-scale sparse functional networks from resting state fMRI. Neuroimage 156, 1–13.

Luciana, M., Bjork, J.M., Nagel, B.J., Barch, D.M., Gonzalez, R., Nixon, S.J., and Banich, M.T. (2018). Adolescent neurocognitive development and impacts of substance use: Overview of the adolescent brain cognitive development (ABCD) baseline neurocognition battery. Dev. Cogn. Neurosci. 32, 67–79.

Lynch, C.J., Breeden, A.L., Gordon, E.M., Cherry, J.B.C., Turkeltaub, P.E., and Vaidya, C.J. (2019). Precision Inhibitory Stimulation of Individual-Specific Cortical Hubs Disrupts Information Processing in Humans. Cereb. Cortex 29, 3912–3921.

Marek, S., Siegel, J.S., Gordon, E.M., Raut, R.V., Gratton, C., Newbold, D.J., Ortega, M., Laumann, T.O., Adeyemo, B., Miller, D.B., et al. (2018). Spatial and Temporal Organization of the Individual Human Cerebellum. Neuron 100, 977–993.e7.

Marek, S., Tervo-Clemmens, B., Nielsen, A.N., Wheelock, M.D., Miller, R.L., Laumann, T.O., Earl, E., Foran, W.W., Cordova, M., Doyle, O., et al. (2019). Identifying reproducible individual differences in childhood functional brain networks: An ABCD study. Dev. Cogn. Neurosci. 40, 100706.

Marek, S., Tervo-Clemmens, B., Calabro, F.J., Montez, D.F., Kay, B.P., Hatoum, A.S., Donohue, M.R., Foran, W., Miller, R.L., Feczko, E., et al. (2020). Towards Reproducible Brain-Wide Association Studies.

Marshall, A.T., Betts, S., Kan, E.C., McConnell, R., Lanphear, B.P., and Sowell, E.R. (2020). Association of lead-exposure risk and family income with childhood brain outcomes. Nat. Med. 26, 91–97.

Mazziotta, J., Toga, A., Evans, A., Fox, P., Lancaster, J., Zilles, K., Woods, R., Paus, T., Simpson, G., Pike, B., et al. (2001). A four-dimensional probabilistic atlas of the human brain. J. Am. Med. Inform. Assoc. 8, 401–430.

Pagliaccio, D., Alqueza, K.L., Marsh, R., and Auerbach, R.P. (2020). Brain Volume Abnormalities in Youth at High Risk for Depression: Adolescent Brain and Cognitive Development Study. J. Am. Acad. Child Adolesc. Psychiatry 59, 1178–1188.

Pauli, W.M., Nili, A.N., and Michael Tyszka, J. (2018). A high-resolution probabilistic in vivo atlas of human subcortical brain nuclei. Scientific Data 5.

Poldrack, R.A., Laumann, T.O., Koyejo, O., Gregory, B., Hover, A., Chen, M.-Y., Gorgolewski, K.J., Luci, J., Joo, S.J., Boyd, R.L., et al. (2015). Long-term neural and physiological phenotyping of a single human. Nat. Commun. 6, 8885.

Power, J.D., Cohen, A.L., Nelson, S.M., Wig, G.S., Barnes, K.A., Church, J.A., Vogel, A.C., Laumann, T.O., Miezin, F.M., Schlaggar, B.L., et al. (2011). Functional Network Organization of the Human Brain. Neuron 72, 665–678.

Power, J.D., Barnes, K.A., Snyder, A.Z., Schlaggar, B.L., and Petersen, S.E. (2012). Spurious but systematic correlations in functional connectivity MRI networks arise from subject motion. Neuroimage 59, 2142–2154.

Power, J.D., Schlaggar, B.L., Lessov-Schlaggar, C.N., and Petersen, S.E. (2013a). Evidence for hubs in human functional brain networks. Neuron 79, 798–813.

Power, J.D., Barnes, K.A., Snyder, A.Z., Schlaggar, B.L., and Petersen, S.E. (2013b). Steps toward optimizing motion artifact removal in functional connectivity MRI; a reply to Carp. Neuroimage 76, 439–441.

Power, J.D., Mitra, A., Laumann, T.O., Snyder, A.Z., Schlaggar, B.L., and Petersen, S.E. (2014). Methods to detect, characterize, and remove motion artifact in resting state fMRI. Neuroimage 84, 320–341.

Rajkowska, G., and Goldman-Rakic, P.S. (1995). Cytoarchitectonic Definition of Prefrontal Areas in the Normal Human Cortex: II. Variability in Locations of Areas 9 and 46 and Relationship to the Talairach Coordinate System. Cereb. Cortex 5, 323–337.

Rosvall, M., and Bergstrom, C.T. (2007). An information-theoretic framework for resolving community structure in complex networks. Proc. Natl. Acad. Sci. U. S. A. 104, 7327–7331.

Rosvall, M., and Bergstrom, C.T. (2008). Maps of random walks on complex networks reveal community structure. Proc. Natl. Acad. Sci. U. S. A. 105, 1118–1123.

Schaefer, A., Kong, R., Gordon, E.M., Laumann, T.O., Zuo, X.-N., Holmes, A.J., Eickhoff, S.B., and Yeo, B.T.T. (2018). Local-Global Parcellation of the Human Cerebral Cortex from Intrinsic Functional Connectivity MRI. Cereb. Cortex 28, 3095–3114.

Scheinost, D., Tokoglu, F., Shen, X., Finn, E.S., Noble, S., Papademetris, X., and Constable, R.T. (2016). Fluctuations in Global Brain Activity Are Associated With Changes in Whole-Brain Connectivity of Functional Networks. IEEE Trans. Biomed. Eng. 63, 2540–2549.

Seitzman, B.A., Gratton, C., Laumann, T.O., Gordon, E.M., Adeyemo, B., Dworetsky, A., Kraus, B.T., Gilmore, A.W., Berg, J.J., Ortega, M., et al. (2019). Trait-like variants in human functional brain networks. Proc. Natl. Acad. Sci. U. S. A. 116, 22851–22861.

Silasi, G., and Murphy, T.H. (2014). Stroke and the connectome: how connectivity guides therapeutic intervention. Neuron 83, 1354–1368.

Sporns, O. (2010). Networks of the Brain (MIT Press).

Sporns, O., Honey, C.J., and Kötter, R. (2007). Identification and classification of hubs in brain networks. PLoS One 2, e1049.

Stein, B.E., and Stanford, T.R. (2008). Multisensory integration: current issues from the perspective of the single neuron. Nat. Rev. Neurosci. 9, 255–266.

Thompson, W.K., Barch, D.M., Bjork, J.M., Gonzalez, R., Nagel, B.J., Nixon, S.J., and Luciana, M. (2019). The structure of cognition in 9 and 10 year-old children and associations with problem behaviors: Findings from the ABCD study’s baseline neurocognitive battery. Dev. Cogn. Neurosci. 36, 100606.

Tustison, N.J., Avants, B.B., Cook, P.A., Zheng, Y., Egan, A., Yushkevich, P.A., and Gee, J.C. (2010). N4ITK: improved N3 bias correction. IEEE Trans. Med. Imaging 29, 1310–1320.

Tyszka, J.M., Michael Tyszka, J., and Pauli, W.M. (2016). In vivo delineation of subdivisions of the human amygdaloid complex in a high-resolution group template. Human Brain Mapping 37, 3979–3998.

Van Essen, D.C., Ugurbil, K., Auerbach, E., Barch, D., Behrens, T.E.J., Bucholz, R., Chang, A., Chen, L., Corbetta, M., Curtiss, S.W., et al. (2012). The Human Connectome Project: a data acquisition perspective. Neuroimage 62, 2222–2231.

Volkow, N.D., Koob, G.F., Croyle, R.T., Bianchi, D.W., Gordon, J.A., Koroshetz, W.J., Pérez-Stable, E.J., Riley, W.T., Bloch, M.H., Conway, K., et al. (2018). The conception of the ABCD study: From substance use to a broad NIH collaboration. Dev. Cogn. Neurosci. 32, 4–7.

Wang, D., Buckner, R.L., Fox, M.D., Holt, D.J., Holmes, A.J., Stoecklein, S., Langs, G., Pan, R., Qian, T., Li, K., et al. (2015). Parcellating cortical functional networks in individuals. Nat. Neurosci. 18, 1853–1860.

Wang, X., Chen, N., Zuo, Z., Xue, R., Jing, L., Yan, Z., Shen, D., and Li, K. (2013). Probabilistic MRI brain anatomical atlases based on 1,000 Chinese subjects. PLoS One 8, e50939.

Weigand, A., Horn, A., Caballero, R., Cooke, D., Stern, A.P., Taylor, S.F., Press, D., Pascual-Leone, A., and Fox, M.D. (2018). Prospective Validation That Subgenual Connectivity Predicts Antidepressant Efficacy of Transcranial Magnetic Stimulation Sites. Biol. Psychiatry 84, 28–37.

Yang, J., and Leskovec, J. (2013). Overlapping community detection at scale. Proceedings of the Sixth ACM International Conference on Web Search and Data Mining - WSDM ’13.

Yeo, B.T.T., Krienen, F.M., Sepulcre, J., Sabuncu, M.R., Lashkari, D., Hollinshead, M., Roffman, J.L., Smoller, J.W., Zöllei, L., Polimeni, J.R., et al. (2011). The organization of the human cerebral cortex estimated by intrinsic functional connectivity. J. Neurophysiol. 106, 1125–1165.

Zhou, Y., Shi, L., Cui, X., Wang, S., and Luo, X. (2016). Functional Connectivity of the Caudal Anterior Cingulate Cortex Is Decreased in Autism. PLoS One 11, e0151879.

