## Supplemental Material for "A Precision Functional Atlas of Network Probabilities and Individual-Specific Network Topography"

SUPPLEMENT

**Supplementary participant information**

Full demographic information for all ABCD Cohorts 1, 2, and 3 are described in Supplementary Table S1, with the exception of participants that were excluded. Of these cohorts, subjects were excluded either because they were unable to be processed through the DCAN processing pipeline (https://github.com/DCAN-Labs/abcd-hcp-pipeline) described in the methods section (typically due to poor image quality) or had fewer than 10 minutes of resting state data post-motion correction.

| Supplementary table 1: Demographics of all ABCD participants for year 1. | | | |
|---|---|---|---|
| **continuous** | **ABCD Cohort-1 (N=5786)** | **ABCD Cohort-2 (N=5786)** | **ABCD Cohort-3 (N=303)** |
| | **mean (sd)** | **mean (sd)** | **mean (sd)** |
| age (months) | 119.01 (7.47) | 118.87 (7.43) | 119.07 (7.81) |
| current grade level | 4.22 (.79) | 4.21 (.79) | 4.28 (.76) |
| highest parent edu. | 17.07 (2.67) | 17.06 (2.66) | 16.75 (2.83) |
| combined income | 7.24 (2.42) | 7.23 (2.42) | 6.93 (2.5) |
| **categorical** | **ABCD Cohort-1 (N=5786)** | **ABCD Cohort-2 (N=5786)** | **ABCD Cohort-3 (N=303)** |
| | **count (%)** | **count (%)** | **count (%)** |
| # female | 2799 (48.4) | 2734 (47.3) | 148 (48.8) |
| anesthesia exposure | 1839 (31.8) | 1828 (31.6) | 87 (28.7) |
| right handed | 4605 (79.6) | 4580 (79.2) | 238 (78.5) |
| **race** | **ABCD Cohort-1 (N=5786)** | **ABCD Cohort-2 (N=5786)** | **ABCD Cohort-3 (N=303)** |
| | **count (%)** | **count (%)** | **count (%)** |
| white | 3719 (64.3) | 3638 (62.9) | 158 (52.1) |
| black | 892 (15.4) | 918 (15.9) | 54 (17.8) |
| AIAK | 27 (.5) | 30 (.5) | 5 (1.7) |

| | | | |
|---|---|---|---|
| NHPI | 10 (.2) | 6 (.1) | 0 |
| asian | 130 (2.2) | 136 (2.4) | 10 (3.3) |
| other | 239 (4.1) | 244 (4.2) | 38 (12.5) |
| unknown/declined | 87 (1.5) | 86 (1.5) | 12 (4.0) |
| more than one race | 682 (11.8) | 728 (12.6) | 23 (7.6) |
| latinx | 1176 (20.6) | 1172 (20.3) | 59 (19.5) |
| **site** | **ABCD Cohort-1 (N=5786)** | **ABCD Cohort-2 (N=5786)** | **ABCD Cohort-3 (N=303)** |
| | **count (%)** | **count (%)** | **count (%)** |
| 1 | 194 (3.4) | 203 (3.5) | 9 (3.0) |
| 2 | 274 (4.7) | 273 (4.7) | 14 (4.6) |
| 3 | 318 (5.5) | 307 (5.3) | 8 (2.6) |
| 4 | 366 (6.3) | 362 (6.3) | 15 (5.0) |
| 5 | 180 (3.1) | 185 (3.2) | 13 (4.3) |
| 6 | 279 (4.8) | 279 (4.8) | 27 (8.9) |
| 7 | 165 (2.9) | 166 (2.9) | 8 (2.6) |
| 8 | 175 (3.0) | 174 (3.0) | 6 (2.0) |
| 9 | 212 (3.7) | 210 (3.6) | 10 (3.3) |
| 10 | 360 (6.2) | 361 (6.2) | 20 (6.6) |
| 11 | 222 (3.8) | 220 (3.8) | 12 (4.0) |
| 12 | 291 (5.0) | 295 (5.1) | 19 (6.3) |
| 13 | 351 (6.1) | 353 (6.1) | 19 (6.3) |
| 14 | 297 (5.1) | 294 (5.1) | 16 (5.3) |
| 15 | 229 (4.0) | 213 (3.7) | 12 (4.0) |
| 16 | 479 (8.3) | 506 (8.7) | 20 (6.6) |
| 17 | 286 (4.9) | 278 (4.8) | 14 (4.6) |
| 18 | 187 (3.2) | 187 (3.2) | 10 (3.3) |
| 19 | 270 (4.7) | 268 (4.6) | 14 (4.6) |
| 20 | 343 (5.9) | 342 (5.9) | 18 (5.9) |
| 21 | 308 (5.3) | 310 (5.4) | 16 (5.3) |

A diagram showing which subjects were used to generate probabilistic maps is shown in Supplementary Figure S1. Brain mapping was performed for all subjects with at least 10 minutes of resting state data using all 3 brain mapping methods: Infomap, template matching, and NMF. We also used template matching and Infomap separately on concatenated rest and task data using the same network templates used for resting state fMRI data.

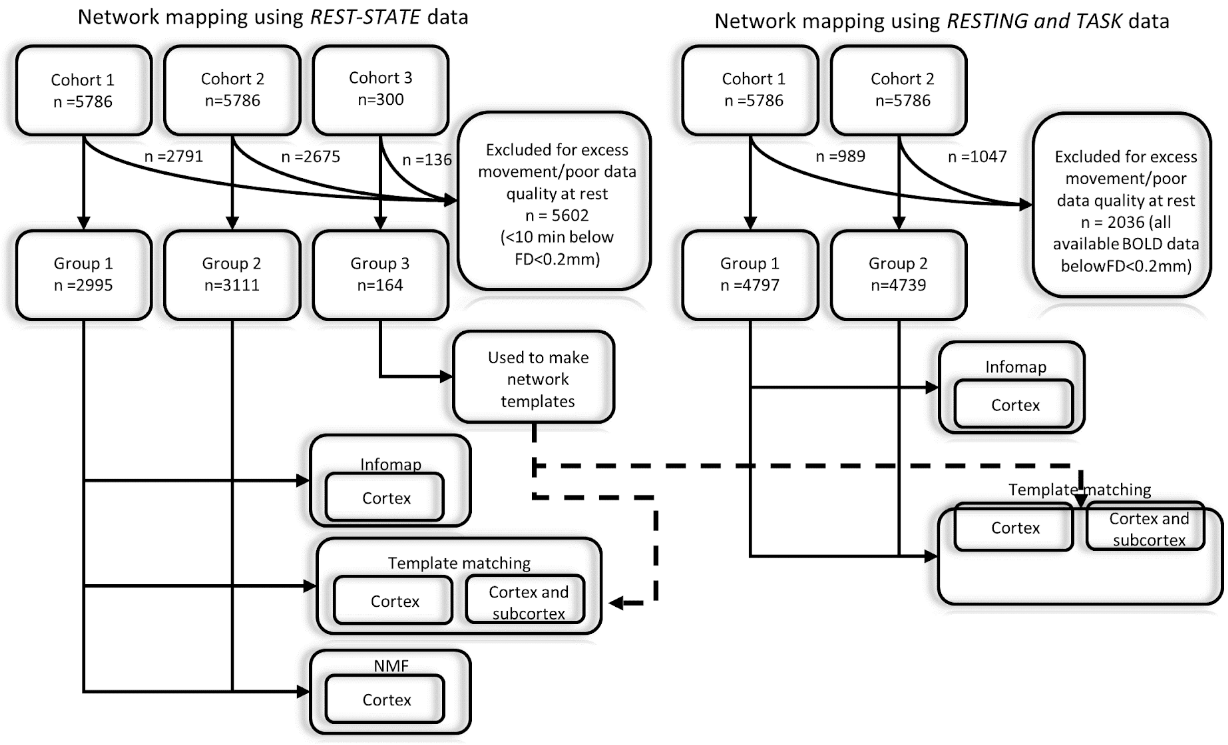

***<u>Supplementary Figure S1)</u>*** *Participant cohorts and the types of community detection implemented. Participants were placed into matched cohorts (see demographics in Supplementary table for additional demographic information of all the subjects).*

**Illustration of Methodologies**

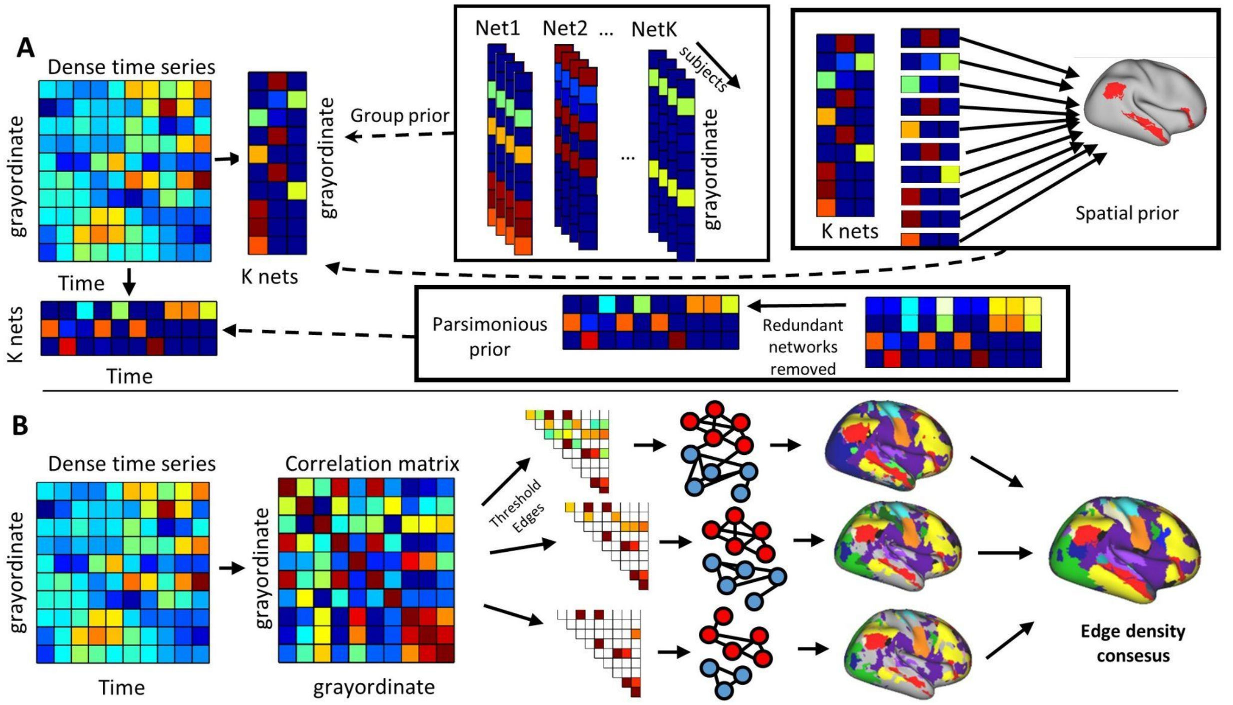

***__Supplementary Figure S2)__ NMF and Infomap methods of community detection.*** *A) Non-negative matrix factorization was conducted on the dense time series using 3 priors: A group consensus prior which encourages subjects to have a similar number of networks, and a spatial prior which serves to constrain the spatial topology of a given network, a parsimonious prior, which removed redundant networks for each subject. B)When Infomap community detection was implemented, we generated a correlation matrix using motion-censored dense time series data (See methods) in an identical manner to the template matching procedure. Each upper triangle of the correlation matrix was then thresholded to various (0.3, 0.4, 0.5, 1.0, 1.5, 2.0, 2.5, 3.0, 3.5, 4.0, 4.5, 5.0) top percentages of the connections. Those connections were then used as the input for Infomap. Infomap uses a random walk to minimize bit-wise code length necessary to describe the whole system structure. The final network labels were then determined by generating a consensus across thresholds, then comparing the Jaccard index of the spatial arrangement of grayordinates from the detected network with those found in the group (See methods).*

In addition to template matching, we also implemented two additional community detection methods: Non-negative matrix factorization (NMF) (Supplementary Figure S2A) and Infomap community detection (Supplementary Figure S2B). Details of both methods are described in the Methods section in the main text. NMF is a group of algorithms in multivariate analysis, where matrix V is the product of two smaller matrices W and H. In Supplementary Figure S2A, NMF decomposes a concatenated dense time series of all the template subjects into 2 constituent factor matrices (a grayordinate by k networks matrix and a grayordinate by time matrix). In contrast to principal component analysis, the components (or networks) are positive additive descriptors of the network.

Similar to how a letter in the U.S. can be addressed to a house with a 2-level description state/province, then city, brain network organization can be described using a 2-level system of networks and nodes respectively.  Infomap is a network-describing algorithm that tries to minimize the number of bits (using Huffman coding) necessary to describe the whole network (Martin Rosvall and Bergstrom 2008; M. Rosvall, Axelsson, and Bergstrom 2009). For example, would it require fewer bits to describe the whole brain with few networks containing many nodes, or many networks with fewer subnodes? Infomap uses a random walk algorithm that uses connection weights to determine the minimum descriptor code length necessary. Importantly, while the solution provides modules, it is not designed to maximize modularity. As others have done previously (Gordon et al. 2017), we thresholded the correlation matrix to the top x% of connections (or edges) because of the computational limitations of using a full set of 8.1 billion connections as descriptors in the map equation.

A graphical description of the template matching method is shown in Supplementary Figure S3. A) Gordon and colleagues (Gordon et al. 2017) generated single network assignments  using Infomap on a group average dense connectivity matrix from a cohort of 210 adults . B) The Gordon parcellation was used to anatomically define networks for each subject and create seed-based correlations for each network in all subjects in the template group (n=164 ABCD Group3  participants). Here we show the default mode network in red as an example. C) Seed-based correlation maps were averaged across the subjects in the template group for each network separately. D) Each template was then thresholded to correlation values >Zscore =1 (~top 15.9% of connections). E) To perform template matching in the group 3 subjects, we first generated whole-brain connectivity matrices. Here, we show an example of the whole-brain connectivity of a grayordinate within the posterior cingulate cortex (PCC). F) We then thresholded the connectivity for each grayordinate in the same manner as the template and calculated an $eta^2$ value for each network. G) Each grayordinate is then assigned the network based on the maximum $eta^2$ value.

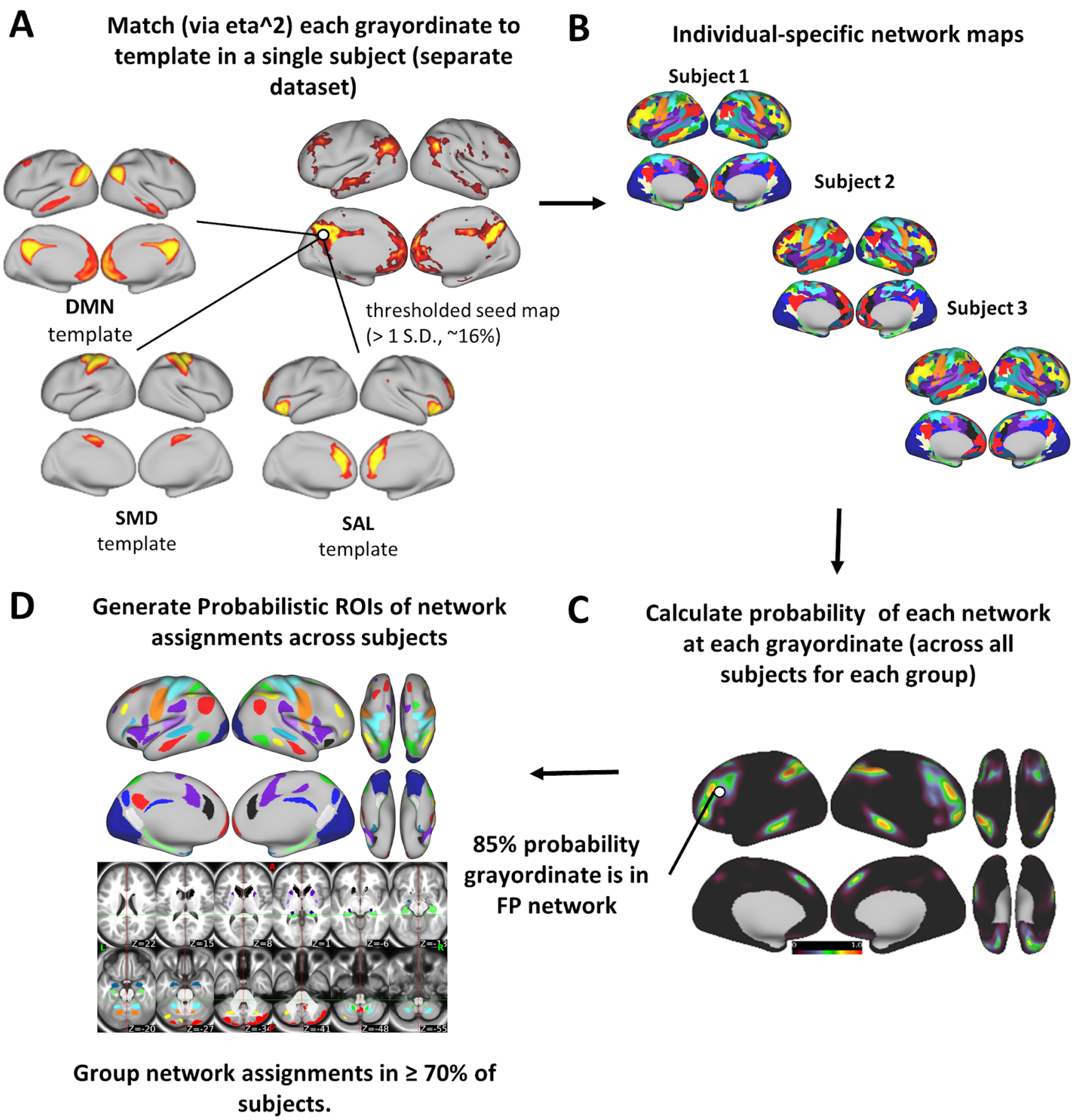

***<u>Supplementary Figure S3)</u> Template matching method.*** *A) For the test subjects, we then generated whole brain connectivity matrices. Each row of the connectivity matrix was compared with each of the network templates. ( only DMN, SMD, and SAL are shown here for visualization purposes). Here, we show an example of connectivity to a grayordinate within the posterior cingulate cortex (PCC), whose connectivity resembles the default mode network. Each template was thresholded to correlation values >Zscore =1 (~top 15.9% of connections). We then thresholded the connectivity for each grayordinate in the same manner as the template and calculated an eta square value for the network. Each grayordinate*

*is then assigned the network based on the maximum eta$^2$ value. B) Each grayordinate was assigned a label for each subject. C) The probability of each network was calculated at each grayordinate. D) We thresholded the probability maps to produce a probabilistic ROI set of high network homogeneity using ABCD group1.*

The network templates generated for template matching are shown in Supplementary Figure S4. Each network was used to compare functional connectivity across experimental groups. The list of networks included are the default mode network (DMN), the visual network (VIS), the frontoparietal network (FPN), the dorsal attention network (DAN), the ventral attention network (VAN), the salience network (Sal), the cingulo-opercular network (CO), the dorsal sensorimotor network (SMd), the lateral sensorimotor network (SMl), the auditory network(AUD), the temporal pole network (TP), the medial temporal network (MTL), the parietal occipital network (PON), and the parietal medial network (PMN).

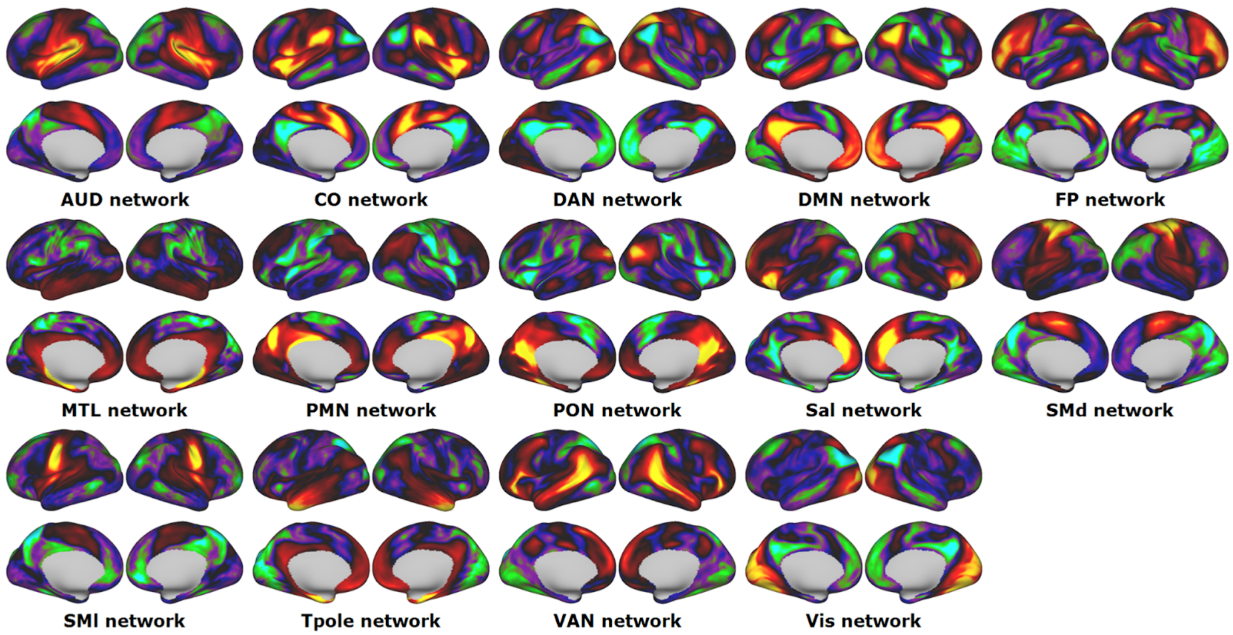

***<u>Supplementary Figure S4)</u>* Network Templates.** *Each network that was used to compare functional connectivity to the experimental groups. The list of networks included are the default mode network (DMN, the visual network (VIS), the frontoparietal network (FPN), the dorsal attention network (DAN), the ventral attention network (VAN), the salience network (Sal), the cingulo-opercular network (CO), the dorsal sensorimotor network (SMd), the lateral sensorimotor Network(SMl), the auditory network(AUD), the temporal pole network (TP), the medial temporal network(MTL), the parietal occipital network(PON), and the parietal medial network(PMN).*

**Reduced data requirements.**

An open question in the field of neuroimaging is: “what amount of resting state data is required to draw reliable conclusions about an individual’s connectome?”. Some estimates examining split-half reliability of connectivity matrices have demonstrated that upwards of 30 minutes of low motion BOLD data is necessary (Laumann et al. 2015). We performed split-half reliability analysis for network maps generated in 10 subjects from the Midnight Scan Club (MSC) data set, who underwent 5 hours of resting state fMRI (in addition to task collection) (Greene et al. 2020; Gordon et al. 2017). We split the resting state scans into interleaved halves, generated networks from each half as described in the Methods, and calculated the NMI between networks generated from halves of the within versus between subjects (identical to the analysis shown in Figure 3B). As with the ABCD dataset, the NMI of networks generated from the same MSC subjects was significantly higher than networks from different subjects. In Supplementary Figure S5, the range of same-subject NMIs is shown in a blue box (0.527-0.648) and the range of null NMI (from comparing different subjects) is shown as a grey box(0.314-0.378). Interestingly, the average intrasubject NMI from MSC subjects was higher than ABCD subjects (MSC: 0.584 vs ABCD:0.4214), suggesting that random sampling from longer/multiple sessions may produce more reliable network maps. In addition to the split halves analysis, we also compared the similarity of networks generated from the second half of a subject’s data (average of 71.28±37.82 min, FD=0.2) vs networks generated from discrete time intervals (1, 2, 3, 4, 5, 10, 15, and 20 min, 10 times each) randomly sampled from the first half (average of 73.12±43.72 min, FD=0.2). The NMI between network maps generated from each interval compared to the second half rapidly increased as correlation matrices contained more time points up to 5 to 10 minutes, then began to plateau. Only 2 minutes of resting state data was needed to generate intrasubject network maps with greater similarity than to the other subjects in the group (1 minutes:$t(9.68)=-3.37$, $p=0.0074$; 2minutes: $t(9.33)=8.919$, $p=7.211 \times 10^{-6}$; 3minutes: $t(9.50)=17.10$, $p=1.858 \times 10^{-8}$; 4minutes: $t(9.26)=15.200$, $p=7.33 \times 10^{-8}$; 5minutes:

t(9.27)=17.554, p=1.978 x $10^{-8}$; 10 minutes: t(9.19)=18.201, p=1.6055 x $10^{-8}$; 15minutes: t(9.20)=20.33, p=5.839 x $10^{-9}$; 20minutes: t(9.15)=18.603, p=1.3864 x $10^{-8}$ ), see Supplementary Figure S5B). For most MSC participants, only 10 minutes of data was required to generate network maps with NMI values that fell within the range of the expected maximum NMI.

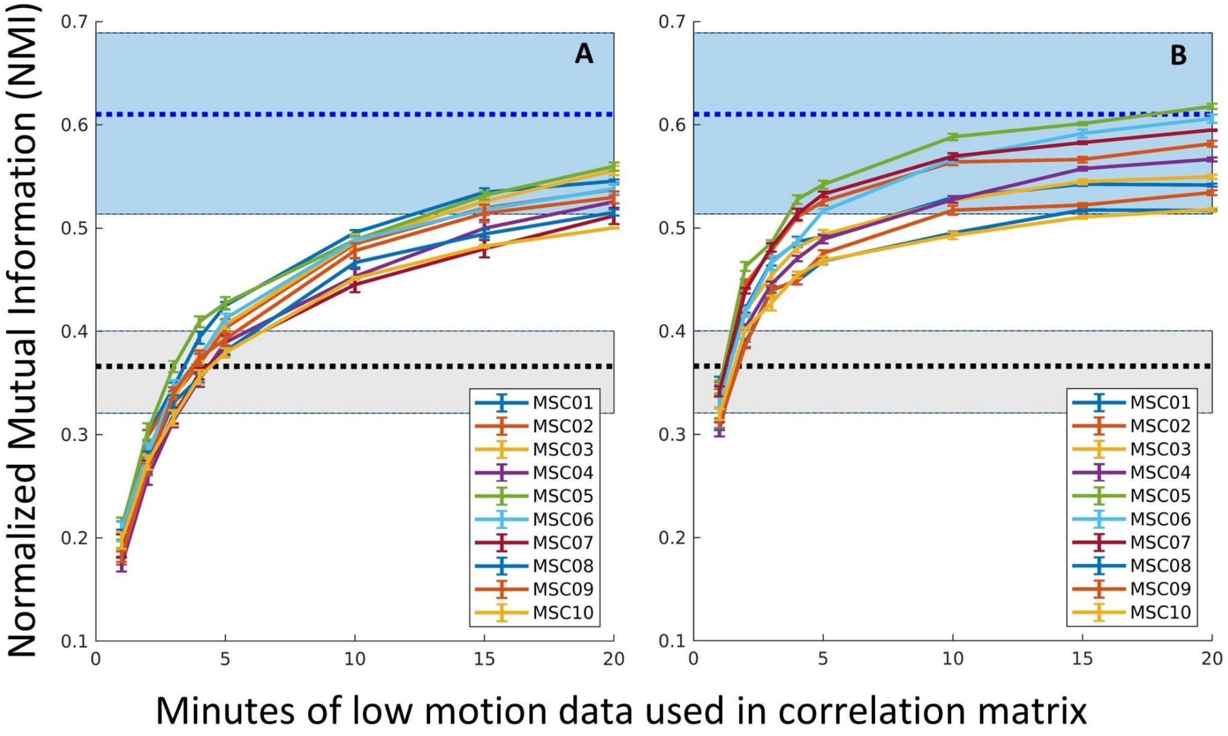

***<u>Supplementary Figure S5:</u>* Calculating networks generated at various time intervals.** *Using data from the Midnight Scan Club (MSC) who underwent 5 hours of resting state fMRI (in addition to task collection)(Sylvester et al. 2020; Greene et al. 2020; Gordon et al. 2017; Gratton et al. 2018). We calculated the NMI between network maps generated from either A) continuous low-motion time frames or B) randomly-sampled low-motion frames, to the participants on hold-out half. The range of the maximum NMI for between participant's own halves using all available motion-censored data is shaded in blue, and the NMI to other participants in the group is shaded in gray. The shaded region indicates the range of the distribution of NMI, using the maximum amount of low motion data for each half, and the thick dotted line represents the mean. Each colored tracing represents 1 subject. Error bars denote ±1 standard error.*

The probabilistic ROIs generated from the ABCD participants used 10 minutes of randomly-sampled data, however sampling from longer data collections, such as the ones collected in the MSC data set, has the potential to artificially inflate the similarity between halves due to the reduced influence of autocorrelation in the timeseries. Therefore, in addition we sampled 10 minutes of continuous low-motion data which was motion censored in an identical manner as described in the methods in the main text, except that time intervals were not randomly sampled throughout the collection, but rather a random low-motion frame was selected, and the amount of following frames corresponding with each time interval were used to generate a correlation matrix. Network maps were then produced by template matching in the same manner as described in the main text. Comparing Supplementary Figure S5A to S5B, we observed that the randomly-sampled frames generated more similar maps between halves (as evidenced by the increase in NMI) compared to the continuously sampled data. The NMI for the group continuously sampled data is significantly greater than the null for time intervals longer than 5 minutes, however, specificity is indicated when data points are no longer in the gray shaded regions(Supplementary Figure S5). All network maps using continuous data for all MSC subjects were outside the gray region after using 10 minutes of continuously sampled data, suggesting that sampling from longer time-intervals does improve reliability which others have shown (Laumann et al. 2015).

### *Network topography observed in the discovery group replicates in a matched sample*

In addition to brain mapping on an individual basis, we also created network maps from average dense connectivity matrices for Groups 1 and 2 to show replication across independent samples and across methods (Supplementary Figure S6). Infomap and template matching brain mapping methods were applied to identical connectivity matrices generated from matched groups. To highlight reproducibility we measured that amount of replication using an *average* dense connectivity matrix generated from all participants within each independent group (Supplementary Figure S6) (Group 1 and Group 2; see Supplementary Table 1 for demographic details). We calculated the NMI between groups and between methods for group-specific networks. The NMI between Group1 and Group2 was relatively high for each method (TM: 0.9110; Infomap: 0.7893), suggesting that each method provides robustness to replication. We also used NMI to compare the similarity of networks generated from TM vs Infomap for each group (Supplementary Figure S6). Between methods, groups generally display similar topographies as evidenced by high NMI values (Group1: 0.4798; Group2: 0.4762), relative to the null comparison between subjects (Figure 1). Infomap (upper row) and template matching (lower row) produced relatively high replication as evidenced by split group NMI. Insets show that the network labels identified for subcortical regions and cerebellum are markedly similar across groups as well.

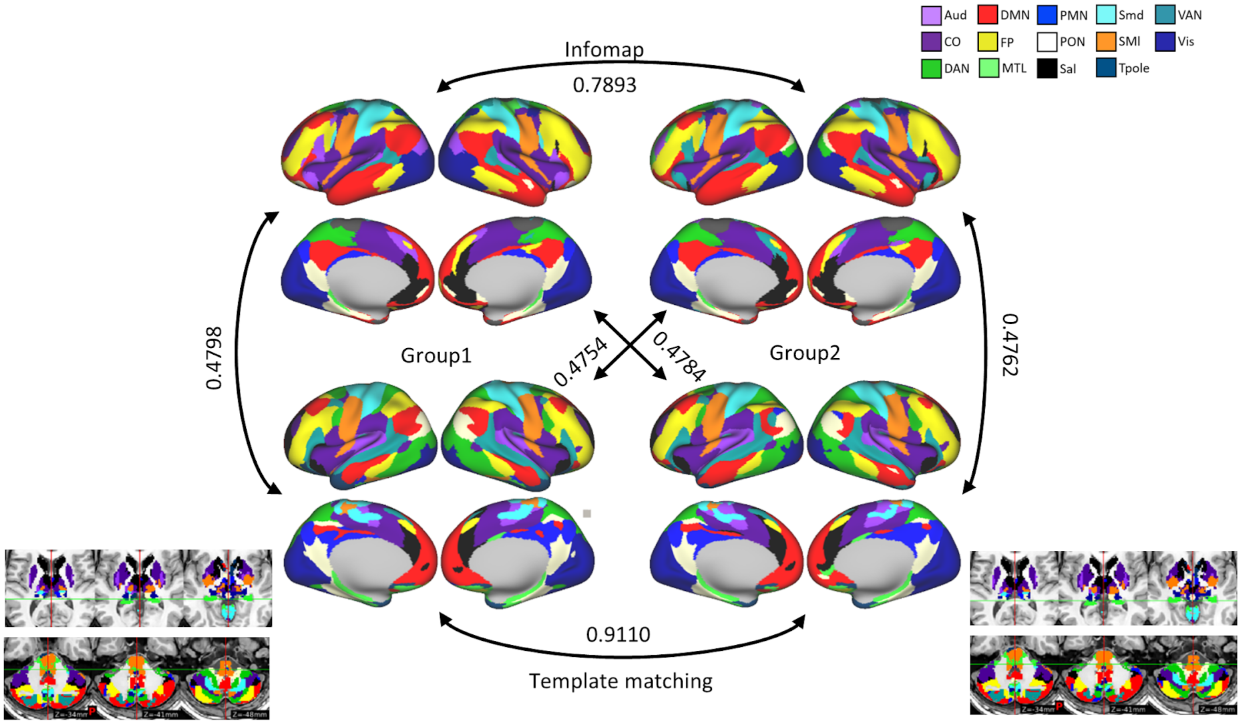

***Supplementary Figure S6: Split half group average comparison with resting state data.*** *Infomap and template matching were run on identical connectivity matrices generated from matched groups. We demonstrate the infomap (upper row) and template matching (lower row) produced relatively high replication as evidenced by split group NMI. Insets: networks labels identified for subcortical regions and cerebellum.*

**Overlapping networks example**

In Supplementary Figure S7A, an example of overlapping networks is shown for an ABCD subject with 10 minutes of low motion resting state data. Because each grayordinate can belong to multiple networks, we used the 10 ABCD subjects mentioned above to calculate NMI independently for each network (Supplementary Figures S7). Networks that have larger topographical variability among the population (e.g. the frontoparietal network) are those that had larger difference in NMI between intrasubject split halves compared to the null distribution, indicating topographical specificity (Supplementary Figure S7B, yellow distribution vs black distribution). Probabilistic mapping of overlapping networks for each group revealed that these probabilistic maps were reliable. (See Figure 6 and Supplementary Figure S11 for all the network maps).

Networks are visualized separately for clarity because each grayordinate can have multiple network assignments. Networks were identified using cortical and subcortical regions. Grayordinates with an $eta^2$ value above the threshold for multiple networks (see Overlapping Template Matching in Methods in main text) received multiple network assignments (see Figure 6 in main text for method). B) For 10 subjects with 20 minutes of low motion data, we performed a similar NMI analysis to that shown in Figure 1 (main text), but we calculated the NMI of split halves for each network separately. Histogram heights have been normalized such that the area under the curve is equal. Note that for networks where topography is highly conserved (such as the lateral somatomotor network), there is considerable overlap of the distributions of NMI of networks for the same vs different subjects. Conversely, the NMI for corresponding halves in networks with highly individualized topology (such as the frontoparietal network) is well outside the null distribution.

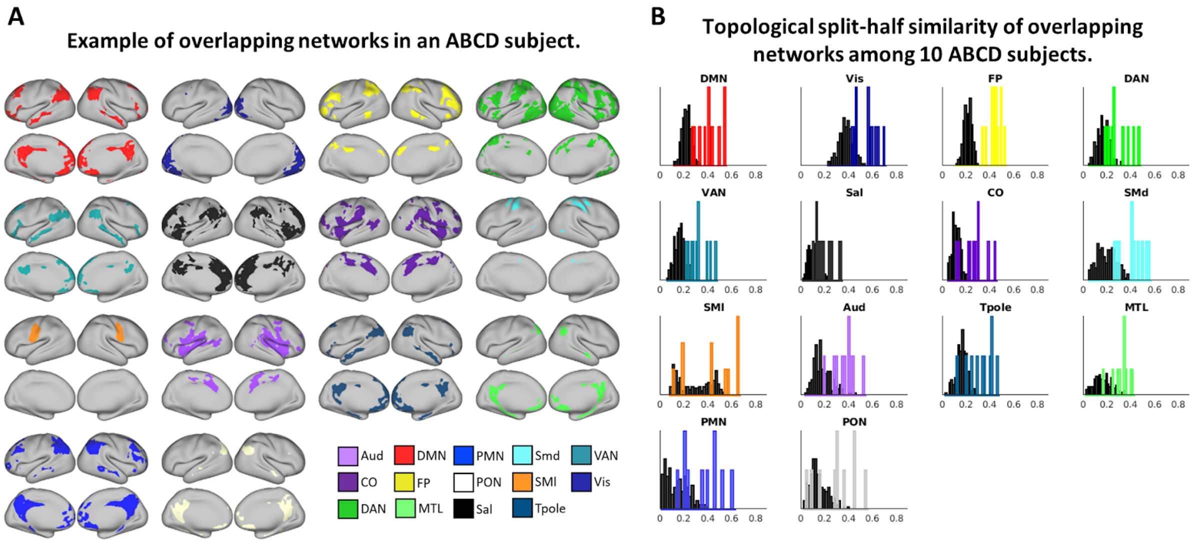

***<u>Supplementary Figure S7</u>*: Overlapping networks from Template matching** *A) Example of overlapping network maps in an ABCD subject. Each network is shown separately for visual comparison. Overlapping Networks are shown for a subject with 10 minutes of low motion resting state data. Networks were identified using cortical and subcortical regions. grayordinates with an eta$^2$ value that is high for multiple networks can therefore receive multiple network assignments (see Figure 6 in main text for method). B) For the 10 subjects 20 minutes of low motion data, we performed a similar NMI analysis to that shown in Figure 3 (main text), but we calculated the NMI for each network separately. Histogram heights have been normalized such that the area under the curve is equal. Note that for networks where topography is highly conserved, the paired NMI heavily overlaps with the NMI for null distribution, however in networks that are highly individualized, the appropriately paired NMI is well outside the null distribution. Colors for networks are identical to those shown in Figure 1.*

**Probabilistic maps**

We showed an example of the probabilistic maps generated by template matching and Infomap algorithms for the frontoparietal cortex in the main text of this manuscript (Figure 2). Here we show the remainder of those maps. Note the strong replication between methods and groups.

**Probabilistic maps – Single network - Template matching: 10 minutes of resting-state data (surface and subcortical)**

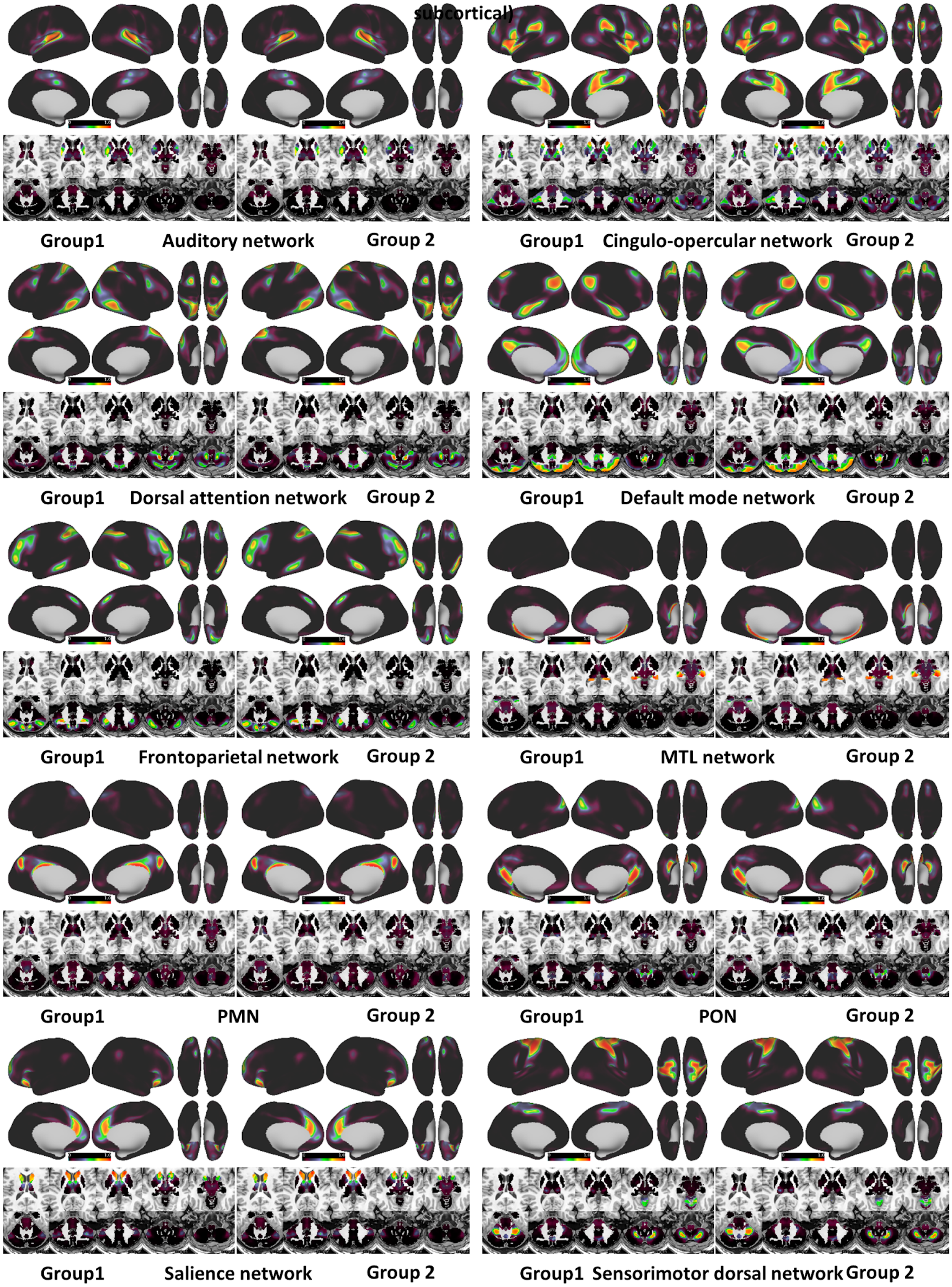

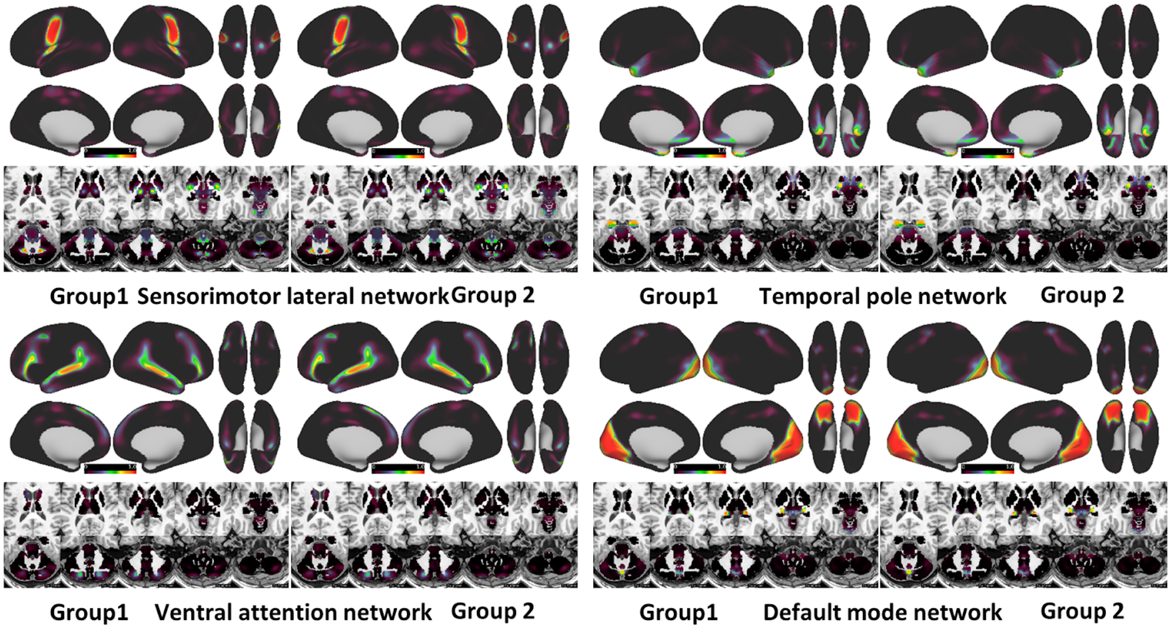

***<u>Supplementary Figure S8</u>*: Probabilistic rest maps (single network)**. *We showed an example of the probabilistic maps generated by template matching and infomap algorithms for the frontoparietal cortex. Here we show the remainder of those maps for template matching. Note the strong replication between methods and groups. The range of all maps is identical (0-1).*

Supplementary Figure S9 shows the same probabilistic network maps as those shown in Figure S8 except that networks were identified using only data from the cortex. This allowed us to compare the topological similarity of our findings to previously identified networks in the literature which have also constrained analysis to the cortex (Gordon et al. 2017)..

Probabilistic maps – single networks - rest using 10 minutes (cortex only)

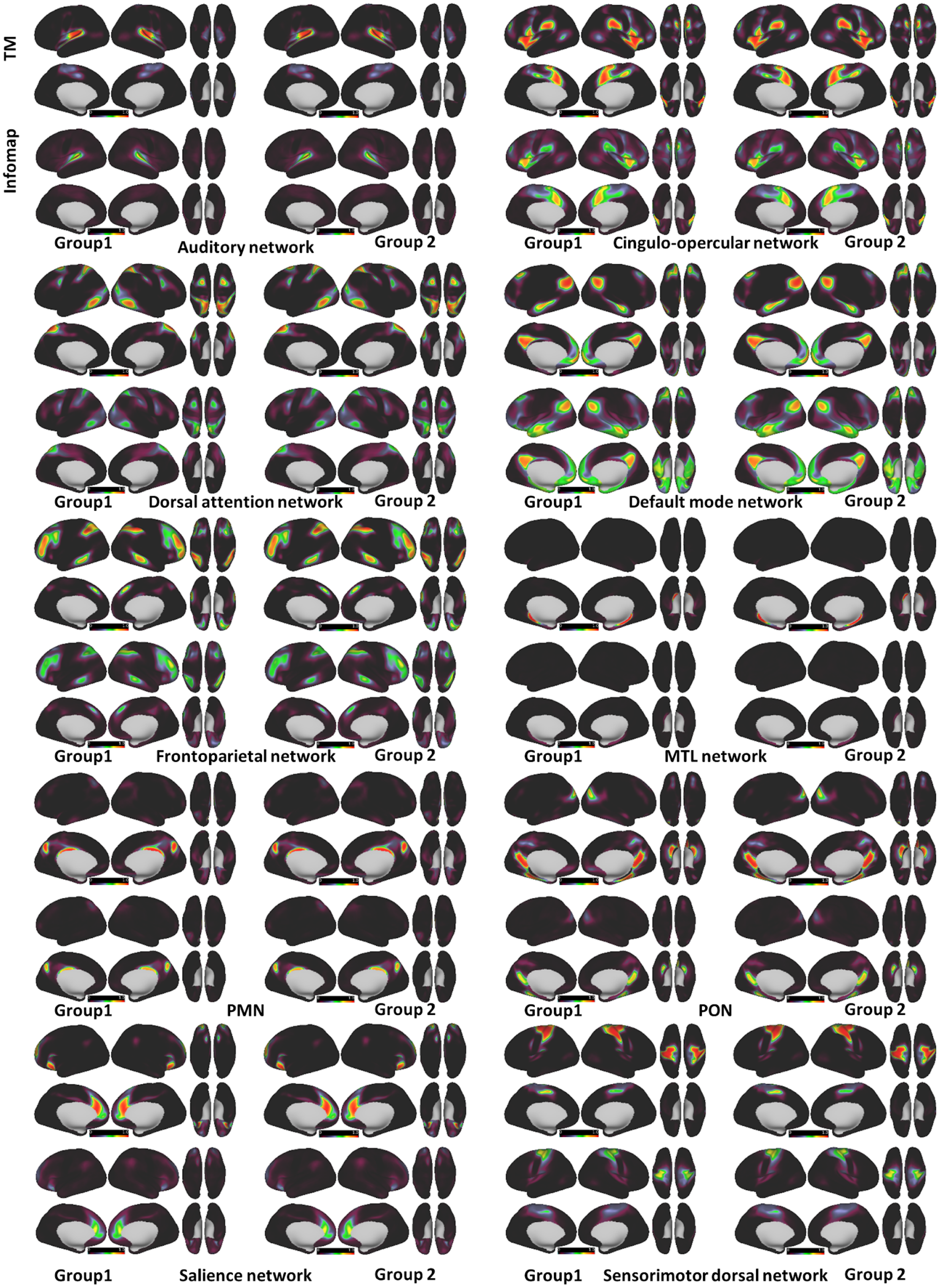

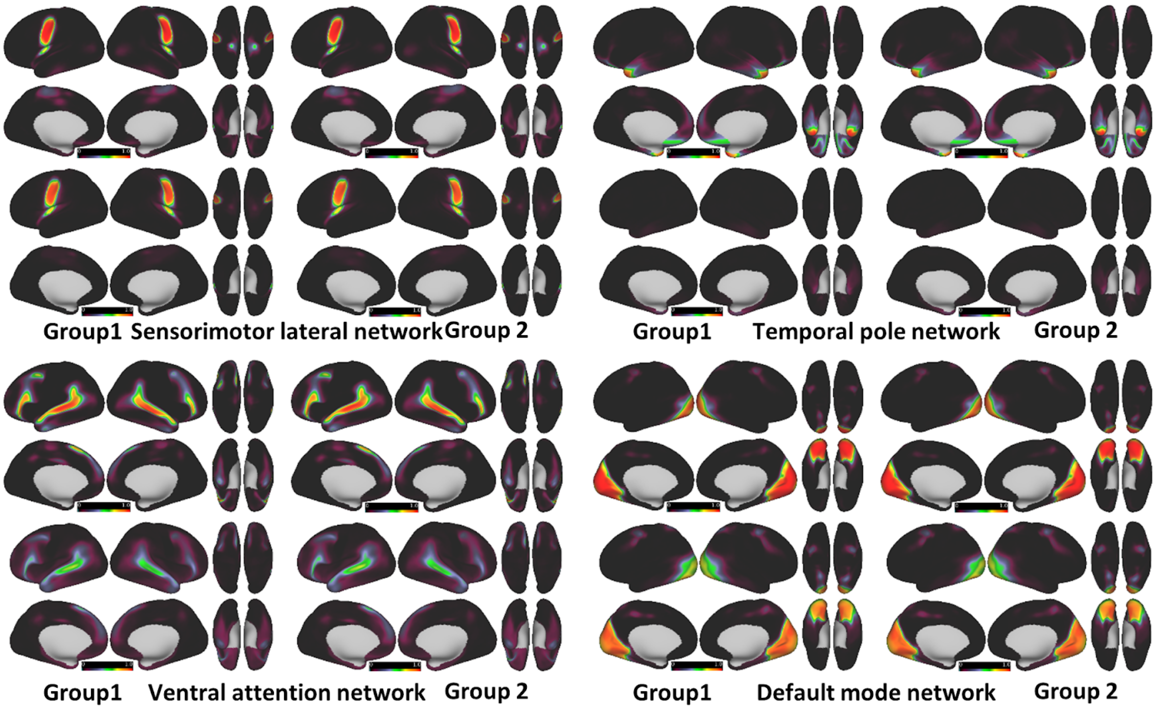

***<u>Supplementary Figure S9:</u>* Probabilistic task maps (single network)***. *The probabilistic maps shown in figure S2 are the same as those shown in figure S1, except that only the correlation data from the cortical surface was used. Infomap community detection was also conducted and is shown here as well (lower images).The range of all maps is identical (0-1).*

In Supplementary Table 2, we've summarized the similarity between groups and methods using only cortical resting state data (as shown in Supplementary Figure S9). We quantified the Pearson correlation between the probabilistic maps that we observed between Groups 1 and 2, and between methods for each group. For any single network, there were a large number of grayordinates with that had zero probability of being labeled as that network. To reduce bias in the intergroup correlation, we only correlated non-zero elements, but we still quantified the number of mismatched zeros (MMZ) to capture subtle differences in topology.

| Supplementary table 2: Correlation between group probabilistic maps using <u>REST</u> data using template matching. *MMZ indicates mismatched zeros. MMZ were excluded from the Pearson correlation shown. | | | |
|---|---|---|---|
| | Template matching | Infomap | Between Methods Single network |

| | Single Net | | Overlapping | | Single Net | | Group1 | | Group2 | |
|---|---|---|---|---|---|---|---|---|---|---|
| | r | %MMZ | r | %MMZ | r | %MMZ | r | %MMZ | r | %MMZ |
| Aud | 0.9997 | 0.119 | 0.9994 | 0.1259 | 0.9994 | 0.0044 | 0.9142 | 0.404 | 0.914 | 0.3794 |
| CO | 0.9997 | 0.0917 | 0.9993 | 0.0986 | 0.9993 | 0 | 0.9184 | 0.228 | 0.9186 | 0.2178 |
| DAN | 0.9997 | 0.1016 | 0.9994 | 0.0921 | 0.9991 | 0.0351 | 0.9468 | 0.2471 | 0.9485 | 0.2641 |
| DMN | 0.9996 | 0.0935 | 0.9994 | 0.1317 | 0.9994 | 0 | 0.7019 | 0.301 | 0.7071 | 0.3042 |
| FP | 0.9996 | 0.0925 | 0.9994 | 0.1339 | 0.9991 | 0.0045 | 0.9613 | 0.2407 | 0.9633 | 0.2295 |
| MTL | 0.9998 | 0.2115 | 0.9999 | 0.2673 | 0.9916 | 0.0452 | 0.8465 | 0.4421 | 0.8511 | 0.4501 |
| PMN | 0.9995 | 0.1045 | 0.9997 | 0.1314 | 0.9994 | 0.025 | 0.9564 | 0.1664 | 0.957 | 0.154 |
| PON | 0.9995 | 0.1147 | 0.9993 | 0.1069 | 0.9991 | 0.0033 | 0.9294 | 0.238 | 0.9299 | 0.2398 |
| Sal | 0.9996 | 0.1458 | 0.9993 | 0.2136 | 0.9988 | 0.0096 | 0.9093 | 0.3808 | 0.9099 | 0.3665 |
| SMd | 0.9998 | 0.139 | 0.9997 | 0.187 | 0.9994 | 0.012 | 0.9564 | 0.3833 | 0.9538 | 0.3356 |
| SMl | 0.9996 | 0.1411 | 0.9998 | 0.0941 | 0.9998 | 0.1669 | 0.9461 | 0.2289 | 0.9441 | 0.2812 |
| Tpole | 0.9996 | 0.1456 | 0.9992 | 0.1372 | 0.9961 | 0.0761 | 0.6792 | 0.2924 | 0.6734 | 0.2719 |
| VAN | 0.9996 | 0.0973 | 0.9994 | 0.1402 | 0.9987 | 0.005 | 0.9446 | 0.2891 | 0.943 | 0.2619 |
| Vis | 0.9999 | 0.1139 | 0.9996 | 0.1095 | 0.9998 | 0.0968 | 0.974 | 0.3566 | 0.9747 | 0.394 |

Probabilistic task maps for template matching (single network) and Infomap using concatenated rest and task data are shown in Supplementary Figure S10. Because global neural network communication appears to be dominated by a shared history of coativation relative to task-induced activations (Gratton et al. 2018), these probabilistic maps are nearly identical (Supplementary Table 3) to those that were observed at rest (Supplementary Figure S9).

**Probabilistic maps – single networks - Template matching TASK (all low motion data –cortex only )**

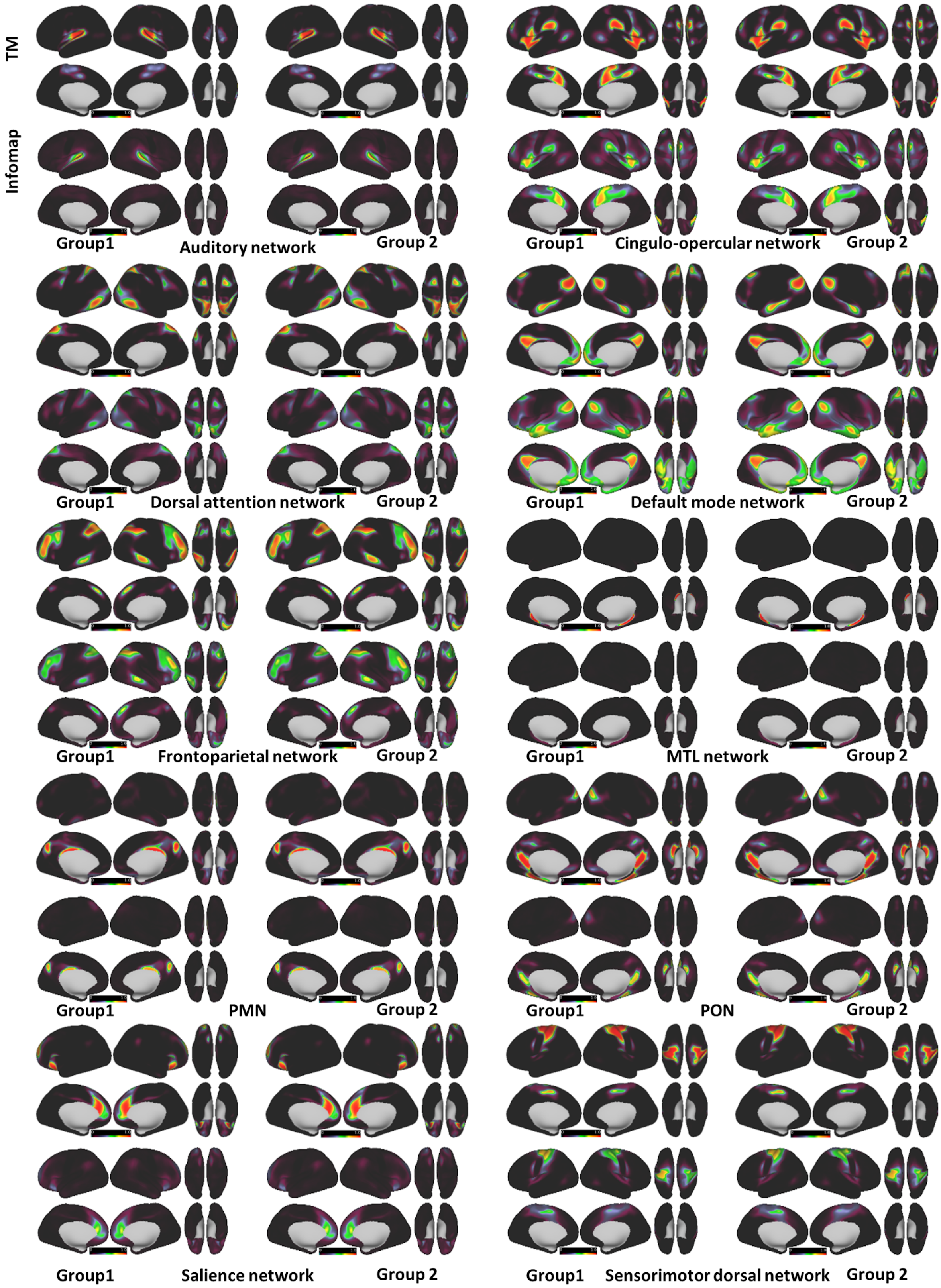

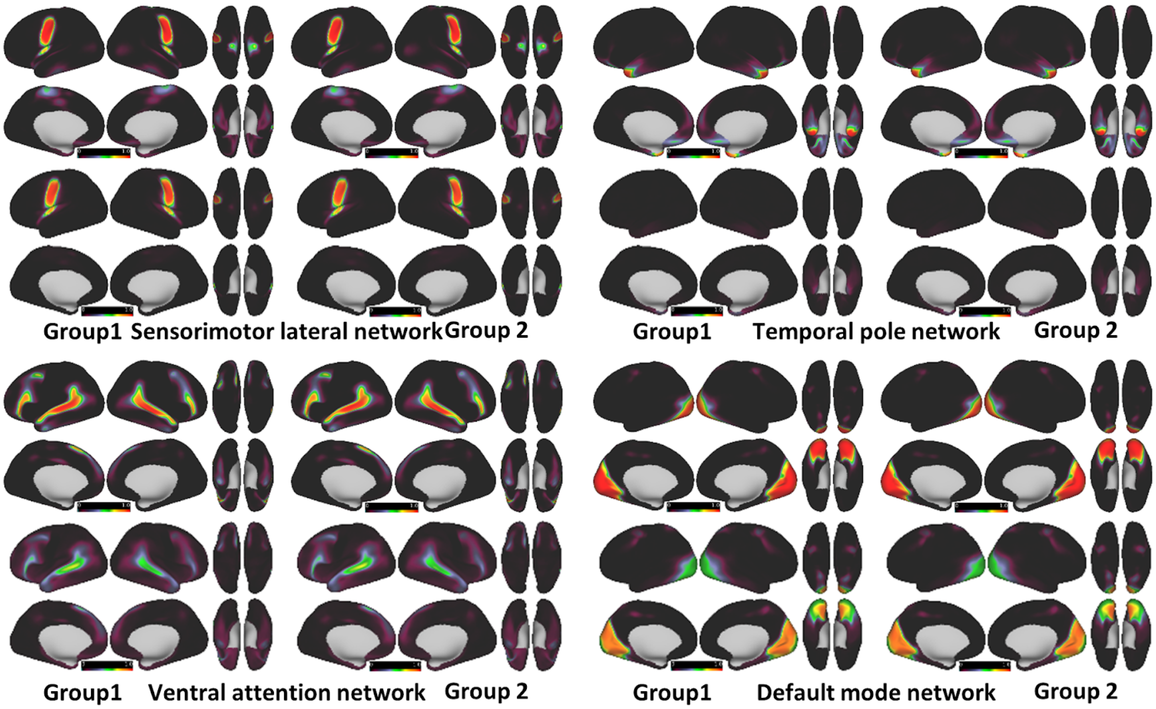

***Supplementary Figure S10: Probabilistic task maps (single network assignment) using template matching.*** *The probabilistic maps shown in figure S2 are the same as those shown in figure S1, except that instead of only using resting state data to generate data, concatenated rest and task data was used.The range of all maps is identical (0-1).*

In Supplementary Table 3, we've summarized the similarity between groups and methods using the concatenated rest and task data (as shown in Supplementary Figure S9) constrained to the cortex, similar to what is shown in Supplementary Table 2. We quantified the Pearson correlation between the probabilistic maps that we observed between Groups 1 and 2, and between methods for each group. Though inter-method correlations are slightly lower compared to rest, between method correlations were still high (0.8769± 0.91 s.d. group1, 0.8769± 0.95 s.d. group2). Less than 1% of grayordinates had mismatched zero probability between groups for any given network.

| Supplementary Table 3: Correlation Between probabilistic maps for each group using TASK data. *MMZ indicates mismatched zeros. MMZ were excluded from the Pearson correlation shown. | | | |
|---|---|---|---|
| | Template matching | Infomap | Between Methods Single network |

| | Single Net | | Overlapping | | Single Net | | Group1 | | Group2 | |
|---|---|---|---|---|---|---|---|---|---|---|
| | r | %MMZ | r | %MMZ | r | %MMZ | r | %MMZ | r | %MMZ |
| Aud | 0.9997 | 0.1224 | 0.9995 | 0.1254 | 0.9993 | 0.0046 | 0.8979 | 0.5251 | 0.8971 | 0.4968 |
| CO | 0.9996 | 0.0911 | 0.9993 | 0.1332 | 0.9993 | 0.0000 | 0.8939 | 0.3247 | 0.8941 | 0.3142 |
| DAN | 0.9996 | 0.1011 | 0.9996 | 0.0988 | 0.9993 | 0.0185 | 0.9371 | 0.3603 | 0.9375 | 0.3553 |
| DMN | 0.9995 | 0.0853 | 0.9994 | 0.0785 | 0.9993 | 0.0001 | 0.6858 | 0.3499 | 0.6754 | 0.3612 |
| FP | 0.9996 | 0.0987 | 0.9994 | 0.1275 | 0.9991 | 0.0025 | 0.9508 | 0.3155 | 0.9538 | 0.2839 |
| MTL | 0.9999 | 0.2075 | 0.9999 | 0.1919 | 0.9885 | 0.0476 | 0.7972 | 0.6120 | 0.7913 | 0.5766 |
| PMN | 0.9994 | 0.1108 | 0.9996 | 0.1265 | 0.9995 | 0.0097 | 0.9159 | 0.2213 | 0.9172 | 0.1995 |
| PON | 0.9995 | 0.1023 | 0.9995 | 0.1128 | 0.9994 | 0.0076 | 0.9025 | 0.3181 | 0.9031 | 0.3082 |
| Sal | 0.9996 | 0.1413 | 0.9994 | 0.2228 | 0.9986 | 0.0061 | 0.8798 | 0.4694 | 0.8895 | 0.4606 |
| SMd | 0.9998 | 0.1419 | 0.9996 | 0.1177 | 0.9995 | 0.0979 | 0.9477 | 0.4556 | 0.9474 | 0.4099 |
| SMl | 0.9996 | 0.1251 | 0.9998 | 0.0846 | 0.9998 | 0.1548 | 0.8819 | 0.2406 | 0.8791 | 0.2518 |
| Tpole | 0.9996 | 0.1635 | 0.9993 | 0.1281 | 0.9958 | 0.0901 | 0.6810 | 0.3975 | 0.6752 | 0.3593 |
| VAN | 0.9997 | 0.0966 | 0.9993 | 0.1796 | 0.9987 | 0.0056 | 0.9500 | 0.3736 | 0.9523 | 0.3283 |
| Vis | 0.9999 | 0.1053 | 0.9996 | 0.1098 | 0.9997 | 0.1096 | 0.9555 | 0.4335 | 0.9557 | 0.3990 |

We generated probabilistic maps with overlapping networks for rest data (Supplementary Figure S11). These maps are similar to the maps shown in Supplementary Figure S8 except that network assignment was allowed to overlap. Note that the region of high probability is generally larger than those shown in Figure S8.

**Probabilistic maps – Template matching Overlapping Networks: 10 minutes of resting-state data**

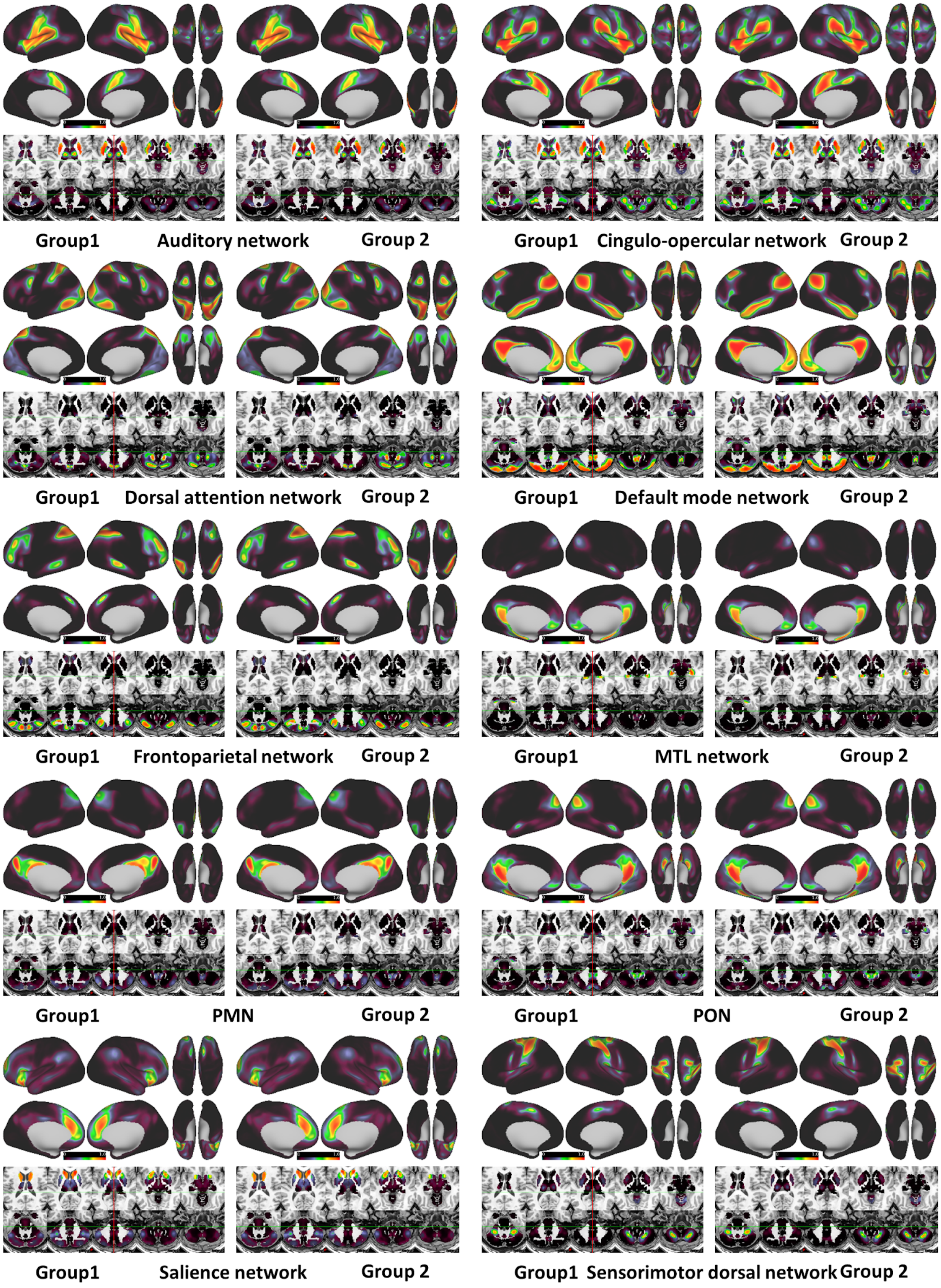

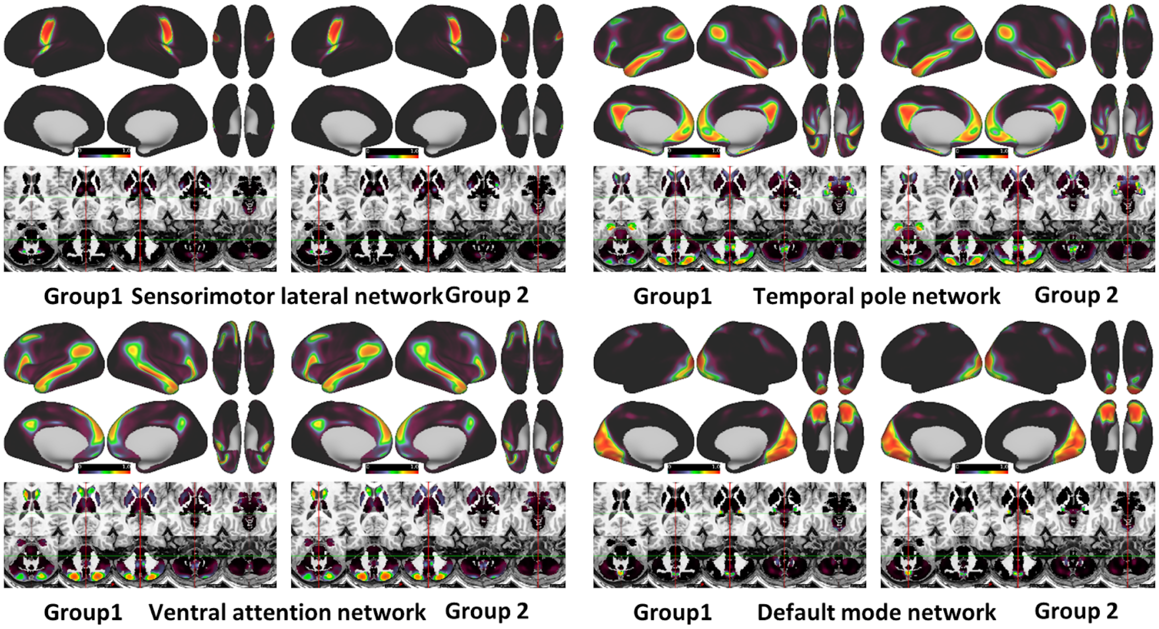

***<u>Supplementary Figure S11</u>*: Probabilistic Rest maps with overlapping networks.** *Network maps were generated using the procedure outlined in figure 5using 10 minutes low-motion of resting state data. The range of all maps is identical (0-1).*

Supplementary Table 4: Correlation Between probabilistic maps from NMF community detection for each group using concatenated <u>TASK</u> data. *MMZ indicates mismatched zeros. MMZ were excluded from the Pearson correlation shown.

| Network | r | %MMZ |
|---|---|---|
| 1 Default Mode Network | 0.99989 | 0.353245 |
| 2 Somatomotor | 0.999897 | 0.004225 |
| 3 Fronto-parietal | 0.999698 | 0.015872 |
| 4 Somatomotor | 0.99984 | 0.003148 |
| 5 Dorsal Attention | 0.999806 | 0.01153 |
| 6 Visual | 0.9999 | 0.003619 |
| 7 Ventral Attention | 0.999587 | 0.00574 |
| 8 Default Mode Network | 0.99971 | 0.009005 |
| 9 Ventral Attention | 0.999718 | 0.013432 |

| | | |
|---|---|---|
| 10 Visual | 0.9999 | 0.002474 |
| 11 Somatomotor | 0.999824 | 0.00409 |
| 12 Default Mode | 0.999711 | 0.011984 |
| 13 Somatoromotor | 0.999899 | 0.003249 |
| 14 Dorsal Attention | 0.999631 | 0.008618 |
| 15 Fronto-parietal | 0.999609 | 0.001818 |
| 16 Auditory | 0.999864 | 0.014054 |
| 17 Frontoparietal | 0.999741 | 0.011227 |

**Volumetric averaging and integration zones**

One might be tempted to interpret the observed integration zones as a byproduct of volumetric averaging due to the limited volumetric resolution of rsFMRI (Supplementary Figure S12D-F). rs-MRI is generally collected with 3-4 mm resolution to optimize the signal-to-noise ratio (SNR) when using a 3 Tesla (3T) scanner (Gorgolewski et al. 2015). There is evidence to suggest that smaller voxels produce a higher SNR and stronger BOLD effects at high fields such as 7T, which, at least within the motor system, can significantly affect the estimate of inter-voxel correlation (Newton et al. 2012). Newton and colleagues demonstrated that BOLD imaging at very high spatial resolution (1×1×2mm) allows for improved functional connectivity analyses, allowing them to distinguish the intricacies of the sensorimotor network (as defined by a finger tapping task compared to rest) in resting state functional connectivity maps. The authors attribute the improvements as partially due to decreased partial volume averaging (Newton et al. 2012). *Any voxel size larger than a single hemodynamic unit (a neuron, corresponding capillaries, and supporting astrocytes) is going to be susceptible to volumetric averaging.* However, while volumetric averaging resulting from our collection resolution (2.4 x 2.4 x 2.4mm) does occur, there are several reasons why it is still likely that neurons residing at the boundaries between networks are important for integration.

First, integration zones appear to be in generally similar locations across the population. If volumetric averaging contributed to the overlapping integration that we've observed, then we would expect them to exist indiscriminately near the boundaries of all networks. Instead, what we observe is that integration zones are present at relatively similar network intersections across participants. Secondly, the location of the integration zones closely corresponds to hubs with regions that are either highly metabolically active (Buckner et al. 2009), relay information between nodes (Sporns 2010), or process multimodal information (Alvarado et al. 2008; Caspers et al. 2006; Vesia and Crawford 2012), which support the hypothesis that that these regions are likely integrating information from multiple networks.

Furthermore, discrete network boundaries such as those shown in Supplementary Figure S12B do not preclude neurons at the interface boundary from communicating with one another. On the contrary, they reinforce the boundary through persistent internetwork communication. Techniques that implement boundary mapping are predicated on the observation that RSFC patterns can abruptly change from one cortical region to an adjacent cortical region, which often reflect the abrupt changes in cytoarchitectonics in the cortex in nonhuman primates (Felleman and Van Essen 1991; Gordon et al. 2016). Few studies have examined cross-network communication at the resolution necessary to capture the nuances of integration between networks. However, in one study of adjacent brain regions, Carmichael and Price (1996) identified two distinct networks within the macaque orbital and medial prefrontal cortex (OMPFC) using retrograde and anterograde tracers (Carmichael and Price 1996). Though the regions networks were clearly distinct, they were highly interconnected at the boundary region between them.(Carmichael and Price 1996; Öngür and Price 2000).

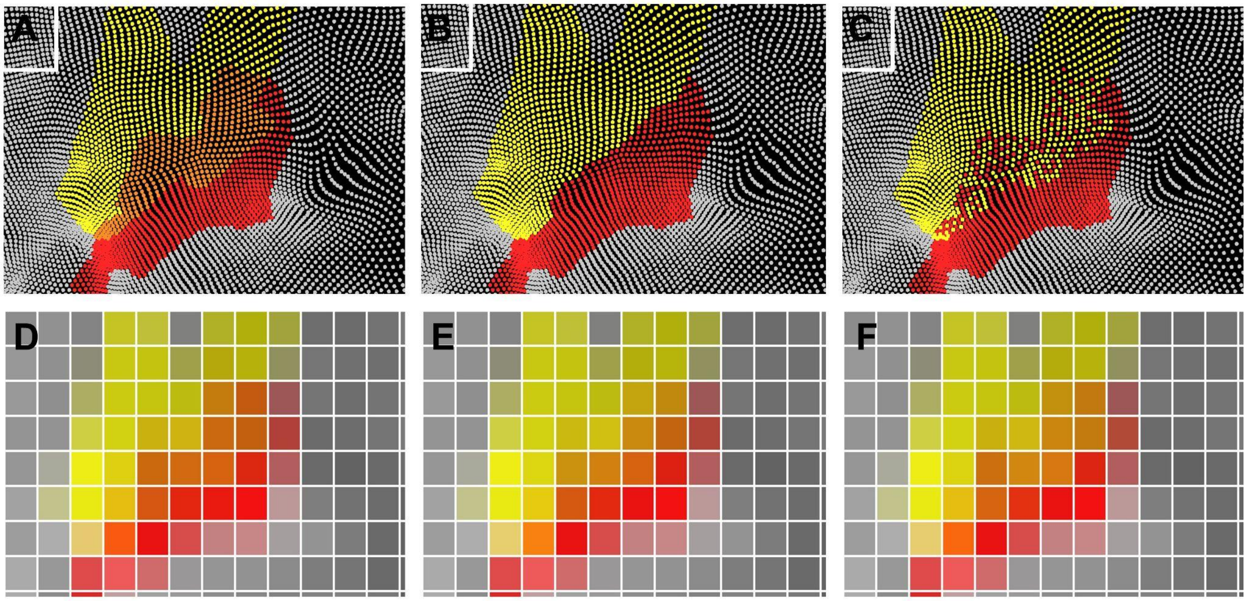

***<u>Supplementary Figure S12</u>*: Schema for how Integration may arise from different topographical arrangements.** *White dots represent a functional unit (e.g. a neuron)* ***A-C)*** *Examples of various topographical border arrangements. Red and yellow represent separate networks. Orange represents functional units that have shared properties of the red and yellow networks. In A, the functional units in orange participate in both the red and yellow networks. In B, functional units establish discrete borders between networks. In C, functional units participate in distinct networks, but are spatially interdigitated.* ***D-F)*** *Volumetric averaging may obscure our ability to distinguish these arrangements (grid not shown to scale for explanatory purposes). Regardless of initial arrangements, functional units that reside at all internetwork boundaries are likely important for integration as well.*

| Supplementary Table 5 of 3-parameter Rise-to-Maximum Fit Coefficient for parcellations shown in Figure 4. | | | | | |
|---|---|---|---|---|---|
| | Coefficient | MIDB Probabilistic Parcellation (80 ROIs) | Gordon Parcellation Full (352 ROIs) | Gordon Parcellation subset (80 ROIs) | Integration Zone Parcellation (30 ROIs) |
| PC1 | y0 | 0.1819±0.0078 | 0.1425±0.0059 | 0.1471±0.0074 | 0.1498±0.0079 |
| | a | 0.5744±0.0088 | 0.5843±0.0071 | 0.5867±0.0090 | 0.6058±0.0100 |
| | b | 0.0031±0.0001 | 0.0033±0.0001 | 0.0032±0.0002 | 0.0031±0.0002 |
| PC2 | y0 | 0.0843±0.0078 | 0.0639±0.0055 | 0.0581±0.0074 | 0.1115±0.0139 |
| | a | 0.4959±0.0149 | 0.4409±0.0110 | 0.4378±0.0132 | 0.5987±0.0225 |
| | b | 0.0022±0.0002 | 0.0021±0.0001 | 0.0022±0.0002 | 0.0024±0.0003 |

| | | | | | |
|---|---|---|---|---|---|
| PC3 | y0 | 0.1239±0.0080 | 0.1034±0.0053 | 0.0979±0.0066 | 0.1254±0.0131 |
| | a | 0.5266±0.0109 | 0.4881±0.0077 | 0.4812±0.0092 | 0.5586±0.0193 |
| | b | 0.0029±0.0002 | 0.0027±0.0001 | 0.0028±0.0002 | 0.0026±0.0003 |

SUPPLEMENTAL REFERENCES

Alvarado, Juan Carlos, Benjamin A. Rowland, Terrence R. Stanford, and Barry E. Stein. 2008. "A Neural Network Model of Multisensory Integration Also Accounts for Unisensory Integration in Superior Colliculus." *Brain Research* 1242 (November): 13–23.

Buckner, R. L., J. Sepulcre, T. Talukdar, F. M. Krienen, H. Liu, T. Hedden, J. R. Andrews-Hanna, R. A. Sperling, and K. A. Johnson. 2009. "Cortical Hubs Revealed by Intrinsic Functional Connectivity: Mapping, Assessment of Stability, and Relation to Alzheimer's Disease." *Journal of Neuroscience*. https://doi.org/10.1523/jneurosci.5062-08.2009.

Carmichael, S. T., and J. L. Price. 1996. "Connectional Networks within the Orbital and Medial Prefrontal Cortex of Macaque Monkeys." *The Journal of Comparative Neurology* 371 (2): 179–207.

Caspers, Svenja, Stefan Geyer, Axel Schleicher, Hartmut Mohlberg, Katrin Amunts, and Karl Zilles. 2006. "The Human Inferior Parietal Cortex: Cytoarchitectonic Parcellation and Interindividual Variability." *NeuroImage* 33 (2): 430–48.

Felleman, D. J., and D. C. Van Essen. 1991. "Distributed Hierarchical Processing in the Primate Cerebral Cortex." *Cerebral Cortex* 1 (1): 1–47.

Gordon, Evan M., Timothy O. Laumann, Babatunde Adeyemo, Jeremy F. Huckins, William M. Kelley, and Steven E. Petersen. 2016. "Generation and Evaluation of a Cortical Area Parcellation from Resting-State Correlations." *Cerebral Cortex* 26 (1): 288–303.

Gordon, Evan M., Timothy O. Laumann, Adrian W. Gilmore, Dillan J. Newbold, Deanna J. Greene, Jeffrey J. Berg, Mario Ortega, et al. 2017. "Precision Functional Mapping of Individual Human Brains." *Neuron* 95 (4): 791–807.e7.

Gorgolewski, Krzysztof J., Natacha Mendes, Domenica Wilfling, Elisabeth Wladimirow, Claudine J. Gauthier, Tyler Bonnen, Florence J. M. Ruby, et al. 2015. "A High Resolution 7-Tesla Resting-State fMRI Test-Retest Dataset with Cognitive and Physiological Measures." *Scientific Data* 2 (January): 140054.

Gratton, Caterina, Timothy O. Laumann, Ashley N. Nielsen, Deanna J. Greene, Evan M. Gordon, Adrian W. Gilmore, Steven M. Nelson, et al. 2018. "Functional Brain Networks Are Dominated by Stable Group and Individual Factors, Not Cognitive or Daily Variation." *Neuron* 98 (2): 439–52.e5.

Greene, Deanna J., Scott Marek, Evan M. Gordon, Joshua S. Siegel, Caterina Gratton, Timothy O. Laumann, Adrian W. Gilmore, et al. 2020. "Integrative and Network-Specific Connectivity of the Basal Ganglia and Thalamus Defined in Individuals." *Neuron* 105 (4): 742–58.e6.

Laumann, Timothy O., Evan M. Gordon, Babatunde Adeyemo, Abraham Z. Snyder, Sung Jun Joo, Mei-Yen Chen, Adrian W. Gilmore, et al. 2015. "Functional System and Areal Organization of a Highly Sampled Individual Human Brain." *Neuron* 87 (3): 657–70.

Newton, Allen T., Baxter P. Rogers, John C. Gore, and Victoria L. Morgan. 2012. "Improving Measurement of Functional Connectivity through Decreasing Partial Volume Effects at 7 T." *NeuroImage* 59 (3):

2511–17.
Öngür, D., and J. L. Price. 2000. “The Organization of Networks within the Orbital and Medial Prefrontal Cortex of Rats, Monkeys and Humans.” *Cerebral Cortex* 10 (3): 206–19.
Rosvall, Martin, and Carl T. Bergstrom. 2008. “Maps of Random Walks on Complex Networks Reveal Community Structure.” *Proceedings of the National Academy of Sciences of the United States of America* 105 (4): 1118–23.
Rosvall, M., D. Axelsson, and C. T. Bergstrom. 2009. “The Map Equation.” *The European Physical Journal. Special Topics* 178 (1): 13–23.
Sporns, Olaf. 2010. *Networks of the Brain*. MIT Press.
Sylvester, Chad M., Qiongru Yu, A. Benjamin Srivastava, Scott Marek, Annie Zheng, Dimitrios Alexopoulos, Christopher D. Smyser, et al. 2020. “Individual-Specific Functional Connectivity of the Amygdala: A Substrate for Precision Psychiatry.” *Proceedings of the National Academy of Sciences of the United States of America* 117 (7): 3808–18.
Vesia, Michael, and J. Douglas Crawford. 2012. “Specialization of Reach Function in Human Posterior Parietal Cortex.” *Experimental Brain Research. Experimentelle Hirnforschung. Experimentation Cerebrale* 221 (1): 1–18.
